# The socio-spatial ecology of giant anteaters in the Brazilian Cerrado

**DOI:** 10.1101/2023.10.04.560744

**Authors:** Aimee Chhen, Alessandra Bertassoni, Arnaud LJ Desbiez, Michael J Noonan

## Abstract

Movement is a key component of an animal’s life history. While there are numerous factors that influence movement, there is an inherent link between a species’ social ecology and its movement ecology. Despite this inherent relationship, the socio-spatial ecology of many species remains unknown, hampering ecological theory and conservation alike. Here, we use fine-scale GPS location data and continuous-time stochastic processes to study the socio-spatial ecology of 23 giant anteaters (*Myrmecophaga tridactyla*) in the Brazilian Cerrado. We found that individuals occupied stable home ranges with a mean area of 5.45 km^2^ with males having significantly larger home ranges than females. The average amount of home-range overlap was low (0.20, n = 121 dyads), with no evidence that giant anteater home ranges were structured based on territorial, mate guarding, nor other social behaviour. We also identified a total of 2774 encounter events. Interestingly, both female-male and male-male dyads had significantly more encounters than female-female dyads, with two pronounced seasonal peaks in female-male encounters. Though encounters occurred frequently, associations between dyads were generally weak and there was little evidence of any correlated movement (mean amount of total correlation = 0.01). Collectively, these findings suggest giant anteaters are a solitary and largely asocial species that readily share space with conspecifics. Despite their present capacity to share space, the combined pressures of being condensed into smaller areas and decreased food availability due to increased pesticide use may see behavioural changes radiating throughout the population. Our study provides insight into heretofore unknown aspects of the socio-spatial ecology of this iconic, but understudied species, as well as crucial information for proactive area-based management. Ultimately, these findings contribute towards sustainable development while potentially maintaining the ecological integrity of giant anteaters and their habitats.

## Introduction

Movement plays a central role in shaping the life histories of motile organisms (Nathan *et al*., 2008; Schick *et al*., 2008; Dougherty *et al*., 2018). An individual’s home range (*sensu* Burt, 1943) will dictate the amount and types of resources it has access to (Kelt and Van Vuren 2001). Their search behaviour will underpin the rate and efficiency with which it encounters those resources (Bartumeus *et al*., 2005; Noonan *et al*., 2023), and the amount of overlap it has with neighbouring individuals will shape where and how often encounters occur (Martinez-Garcia *et al*., 2020; Noonan *et al*., 2021). Movement thus represents an important mechanistic link between individual behaviour and many higher-level ecological processes (Nathan *et al*., 2008; Schick *et al*., 2008; Dougherty *et al*., 2018). In turn, the factors that shape an individual’s movement can have profound impacts on its life history (Leblond, Dussault & Ouellet, 2013; Shiratsuru *et al*., 2023). Understanding species’ movement ecology is thus not only valuable from a natural history perspective, but essential for conservation efforts in protecting species and managing populations (Allen & Singh, 2016; Fraser *et al*., 2018; Bro-Jørgensen, Franks & Meise, 2019) especially in behavioural response to human-activity such as anthropogenic development (Whittington *et al*., 2022).

While there are numerous factors that will influence an animal’s movement (*sensu,* Nathan *et al*., 2008), there is an inherent link between a species’ social ecology and its movement ecology (Hertel *et al*., 2020; Webber *et al*., 2023). For example, in species that maintain actively defended territories (Powell, 2000), interactions between neighbouring animals can constrain movement and limit access to resources (Moorcroft, Lewis & Crabtree, 1999; Tórrez-Herrera, Davis & Crofoot, 2020). In group-living species, shared knowledge and decision-making can lead to collective motion (Strandburg-Peshkin *et al*., 2015) and/or more efficient movement (Mueller *et al*., 2013; Jesmer *et al*., 2018). Similarly, the distribution and abundance of resources will influence their defensibility (*sensu* Grant, 1993) and the propensity towards group-living (Macdonald, 1983; Macdonald & Johnson, 2015). A species’ movement ecology is thus tightly intertwined with its social behaviour (Webber *et al*., 2023). Yet, even with rapid technological (Kays *et al*., 2015; Nathan *et al*., 2022) and analytical (Calabrese, Fleming & Gurarie, 2016; Calabrese *et al*., 2018; Winner *et al*., 2018) advances, there are many species for which details on the socio-spatial interface remain unknown and these knowledge gaps present a notable conservation challenge.

Giant anteaters (*Myrmecophaga tridactyla*) are a prime example of a species for which there is a limited understanding of their socio-spatial ecology. Native to Central and South America (Miranda, Bertassoni & Abba, 2014), giant anteaters are generally found in open grasslands and forest habitats (Mourão & Medri, 2007; Di Blanco, Jiménez Pérez & Di Bitetti, 2015). They have been experiencing population declines due to habitat loss, vehicle collisions, and fires (Miranda *et al*., 2014; Ascensão & Desbiez, 2022; Noonan *et al*., 2022), and are currently classified by the IUCN as vulnerable to extinction, with a risk of local extirpations throughout their range (Miranda *et al*., 2014). Their lifespan in the wild is currently unknown, they have a low reproductive rate (Miranda *et al*., 2014; Desbiez, Bertassoni & Traylor-Holzer, 2020) and occur at low densities (<1/km^2^) (Bertassoni, Bianchi & Desbiez, 2021). Ongoing habitat loss throughout their range (Strassburg *et al*., 2017) is expected to further threaten their survival. Yet, due to large gaps in our knowledge of their natural history, the way in which giant anteater populations will (re)distribute themselves in these changing, fragmented landscapes is unknown.

Here, we address this research gap and focused on answering two overarching questions: i) is the spatial arrangement of giant anteater home ranges governed by social factors (e.g., territoriality, mating groups, social groups, etc.)?; and ii) does giant anteater movement within their home range exhibit signs of social interactions (e.g., avoidance, attraction, correlated movement, etc.)? Currently, little is known about the socio-spatial ecology of giant anteaters (Bertassoni & Desbiez, 2022). Existing studies on free-ranging individuals have shown that home ranges can overlap within and between sexes (Bertassoni & Ribeiro, 2019; Bertassoni, Mourão & Bianchi, 2020), a few instances of mating and courtship behaviour have also been previously recorded (Shaw, Machado-Neto & Carter, 1987; Júnior & Bertassoni, 2014), however, these relationships are still poorly understood. In addition, encounters involving intraspecific interactions such as meeting, following, and foraging together are known to occur (Bertassoni & Milléo Costa, 2010; Catapani *et al*., 2020). As such, we expected to observe a high amount of overlap between individuals with the potential for some amount of correlated movement. Likewise, because of the lack of a clearly defined mating season (Bertassoni & Desbiez, 2022), encounters were expected to be infrequent without any seasonal patterns. Findings are directly applicable for sustainable development as an integrated approach towards area-based management for the conservation of giant anteaters.

## Materials and Methods

### Data collection

The data used in the present study were collected as part of a broader giant anteater monitoring program being carried out in the Cerrado, a savannah-like biome within the Mato Grosso do Sul (MS) state of Brazil. The landscape was primarily dominated by pastures and eucalyptus plantations, interspersed with fragmented natural forest and savanna habitats (Souza *et al*., 2020). Between 2017 and 2018 wild giant anteaters were captured and equipped with GPS tracking harnesses. Full details on the capture and handling protocol are provided in (Kluyber *et al*., 2021), but an overview is provided here. A team comprised at minimum of two veterinarians and a wildlife biologist located and captured giant anteaters in open areas using two long-handled dip nets. Immobilised individuals were then anaesthetized using an intramuscular anesthetic injection comprised of butorphanol tartrate (0.1 mg kg^-1^, detomidine hydrochloride (0.1 mg kg^-1^), and midazolam hydrochloride (0.2 mg kg^-1^) administered to the hind limbs. During immobilization, physical exams were conducted to evaluate health conditions, and individuals deemed healthy were fitted with a GPS tracking harness (TGW-4570-4 Iridium GPS and VHF transmitters (MOD 400; Telonics, Mesa, Arizona).

The GPS harnesses were programmed to record GPS locations at 20-minute intervals. Individuals were recaptured ca. 1 year later for harness removal and the data were downloaded. Every two weeks during the tracking period, each giant anteater was monitored, and health was visually inspected from a distance to minimize disturbance. While a total of 43 individuals were collared as part of the larger monitoring effort (Noonan *et al*., 2022), here we restricted our analyses to 23 range-resident individuals living in three separate clusters (Fig. 1). These individuals were selected as they resided in areas where there was high confidence that all resident giant anteaters were equipped with GPS trackers. The resulting dataset consisted of 528,324 GPS fixes.

**Figure 1.**
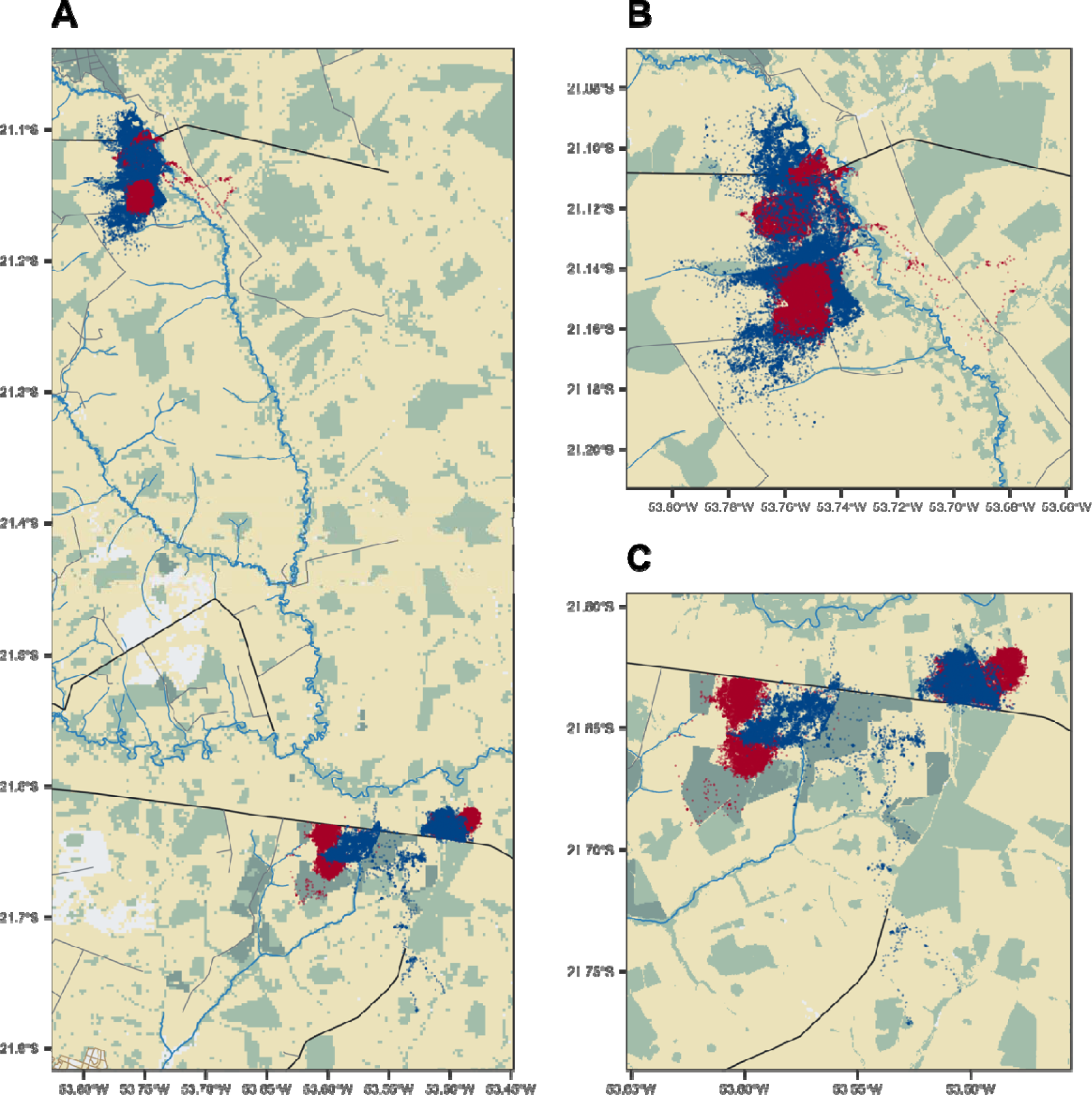
Map of giant anteater location data. Scatter plot depicting the GPS derived location data for 23 giant anteaters in the Brazilian Cerrado, blue points indicate males and red females. The colours in the background depict the landcover classifications, with yellow showing pasture, light green planted forest, and dark green native forest. obtained from MapBiomas. Roads and waterways are also depicted in black and blue respectively.

### Data analysis

Data visualisation and analysis were carried out using R (version 4.2.2, R Core Team, 2022) and the following R packages: `ctmm` (version 1.1.0, Fleming & Calabrese, 2022), lme4 (version 1.1.31, Bates *et al*., 2015), `glmmTMB` (version 1.1.7, Brooks *et al*., 2017), `corrMove` (version 0.1.0, Calabrese & Fleming, 2023), and `ggplot2` (version 3.4.2, Wickham, 2016). All R scripts can be found in the GitHub repository at https://github.com/QuantitativeEcologyLab/Giant_Anteater. Prior to conducting the analyses described below, we calibrated the GPS measurement error and removed any outlying data points (for full details on data pre-processing see Appendix S2 in Noonan *et al*., 2022). In order to answer our two core research questions, we carried out three separate analyses focused on estimating patterns in i) home-range overlap; ii) proximity and encounter rates; and iii) correlated movement. The specific steps carried out in each of these analyses are described below.

#### Home-range overlap

We estimated giant anteater home ranges using Autocorrelated Kernel Density Estimation (AKDE; Fleming *et al*., 2015) using the `ctmm` package. AKDE, corrects for autocorrelation induced bias (Noonan *et al*., 2019) by conditioning the bandwidth optimisation on the data’s autocorrelation structure (Fleming & Calabrese, 2017). Thus, home-range estimation first required fitting a series of continuous-time movement models to the GPS tracking data and identifying the best model via small sample-sized corrected Akaike’s Information Criterion (AIC_c_) (Fleming & Calabrese, 2022). Giant anteater home ranges were then estimated conditional on each individual’s best-fit movement model. Finally, home-range overlap was estimated for all pairs of individuals (i.e., dyads) via the Bhattacharyya coefficient (Winner *et al*., 2018). To determine if sex was a factor underpinning the degree of pairwise overlap, we fit a Generalized Linear Mixed Model (GLMM) with a beta distribution and a logit link function to the home-range overlap estimates, with pairwise sex as a predictor variable (i.e., male-male, female-female, and female-male). In addition, site was included as a random effect. Because the overlap values ranged between [0,1], we rescaled the values as (y(n-1) + 0.5)/n in order for them to lie within the (0,1) interval (Smithson & Verkuilen, 2006). This model was then compared to a similar model that excluded the pairwise sex predictor variable using a likelihood ratio test.

#### Proximity and encounter rates

While home-range overlap describes patterns in spatial structuring, it does not directly indicate whether individuals are likely to be in the same place at the same time (Winner *et al*., 2018). In order to understand the temporal component of giant anteaters’ socio-spatial behaviour, we estimated a proximity ratio for all dyads via the `ctmm` function `proximity()`. The proximity ratio was estimated by comparing a dyad’s observed separation distances with the separation distances expected under completely random movement. A proximity ratio of 1 is thus consistent with independent movement, values <1 indicate that the two individuals are closer on average than expected for independent movement, and vice versa for values >1. We used a GLMM with a gamma distribution, a log link function, and site as a random effect to determine whether the estimated proximity ratios differed between pairwise sex combinations. This model was then compared to similar model that excluded the pairwise sex predictor variable using a likelihood ratio test.

We also estimated the Euclidean distance between the individuals in each dyads at each timepoint using the `ctmm` function `distances()`. From these separation distances we estimated the number of encounter events for each dyad using a 15m distance threshold which, from a behavioural perspective for the species, indicates that individuals are in close enough proximity for an encounter to have occurred. A sensitivity analysis on the 15m threshold is provided in Appendix S1. We used a GLMM with a Poisson distribution, a log link function, and site as a random effect to determine whether encounter rates differed between the pairwise sex combinations. This model was then compared to similar model that excluded the pairwise sex predictor variable using a likelihood ratio test. Lastly, we used a Hierarchical Generalized Additive Model (HGAM) with a Poisson distribution, a log link function, to test for any temporal trends in encounters.

#### Correlated movement

To evaluate if pairs of giant anteaters exhibited any correlation in their movement, all dyads with a proximity ratio that differed significantly from 1 (based on the 95% confidence interval) were identified and carried forward to our subsequent, correlated movement analysis. For these 12 dyads, we used the methods implemented in the `corrMove` package to estimate the amount of correlation in the drift and diffusion components of those dyads’ movement (Calabrese *et al*., 2018). Under an assumption of Brownian motion, the `corrMove` algorithm estimates transition points between a family of four possible movement models; uncorrelated drift and uncorrelated diffusion (UU), correlated drift and uncorrelated diffusion (CU), uncorrelated drift and correlated diffusion (UC), and correlated drift and correlated diffusion (CC) (Calabrese *et al*., 2018). The most appropriate model for any time window was identified via AICc based model selection. From these analyses, we obtained information on the length of time for which the dyad exhibited behaviour consistent with the selected movement model, as well as the correlation coefficients for the drift and diffusion terms (ranging between –1, indicating avoidance, and 1, indicating following).

## Results

### Giant anteater home range arrangement

The 23 range-resident individuals we monitored occupied stable home ranges with a mean area of 5.45 km^2^ (95% CI: 4.55 – 6.45 km^2^; Fig. 1A). We found that males had slightly larger home ranges than females (male 7.28 km^2^, 95% CI: 3.69 – 12.94 km^2^; female 3.32, 95% CI: 2.56 – 4.23 km^2^). We estimated the female/male ratio of mean home-range areas to be 0.41 (0.19–0.85), which excludes 1, indicating statistical significance. Overall, we found no evidence that giant anteater home ranges were structured based on territorial, mate guarding, nor other social behaviour (Fig. 2C, 2D). Of the 121 unique dyads we worked with, the average amount of home-range overlap was 0.20 (range: 0.00 - 0.96). The mean home-range overlap was 0.22 (range: 0.00 – 0.87) for the 31 male-male dyads, 0.14 (range: 0.00 – 0.87) for the 25 female-female dyads, and 0.21 (range: 0.00 - 0.96) for the 65 female-male dyads. Though female-female dyads tended to have the lowest overlap (Fig. 2B), a likelihood ratio test found no evidence that sex impacted the amount of home-range overlap (p = 0.34).

**Figure 2.**
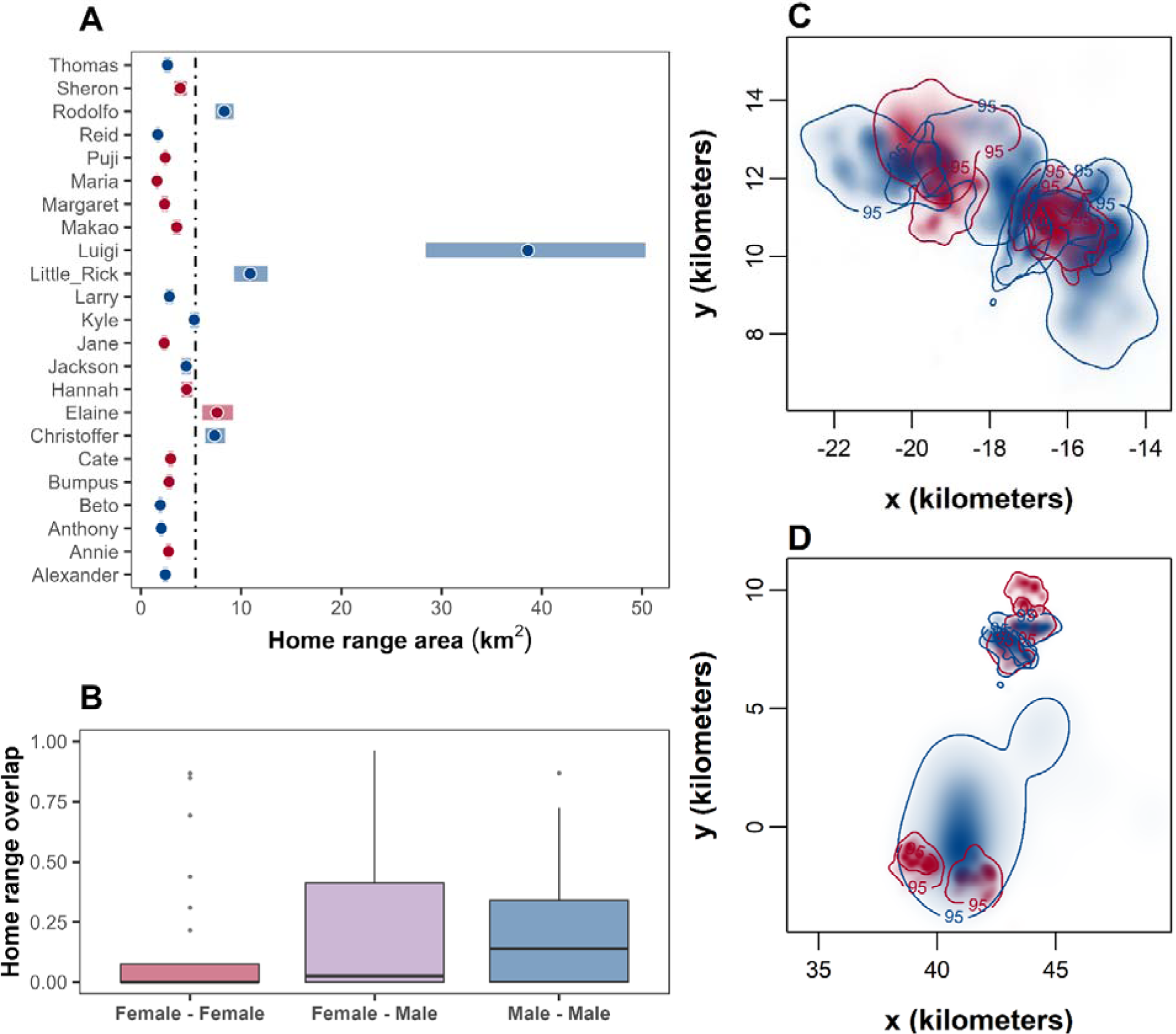
Trends in giant anteater home-range overlap. Panel A) shows a scatter plot of home-range size of each giant anteater. The width of each bar indicates the 95% confidence intervals on the individual home-range estimates. The dashed line denotes the mean home-range size. The boxplots in B) depict the distribution of the home-range overlap estimates between the different combinations of sexes (n = 121). Panels C) and D) show the estimated home-range probability density functions and 95% contours. In all panels, blue indicates male, red indicates female, and purple indicates female-male.

### Giant anteater encounters and inter-individual associations

From the estimated proximity ratios, we found that the associations between dyads were generally weak. Of the 121 dyads, only 12 had both non-zero overlap and proximity ratios that differed significantly from the expectations under independent movement (Fig. 3A). Of these 12 dyads, one male-male pair had a proximity ratio above 1, indicating avoidance behaviour. The remaining 11 dyads had proximity ratios below 1, indicating that these dyads were closer on average than expected for independent movement. These 11 dyads included 2 male-male pairs, 3 female-female pairs and 6 female-male pairs. From a likelihood ratio test, there was no evidence that the estimated proximity ratio differed between sexes for all 121 dyads (p = 0.13) nor for the 12 identified dyads with a non-zero overlap and proximity ratios (0.16), though we did find that there was a weak but significant negative relationship between the amount of home-range overlap and the pairwise proximity ratio (β = −0.12 ± 0.056 standard error (SE), p = 0.025, Fig. 3A) across all 121 dyads but not for the 12 identified dyads with a non-zero overlap and proximity ratios (β = 0.16 ± 0.349 standard error (SE), p = 0.644).

**Figure 3.**
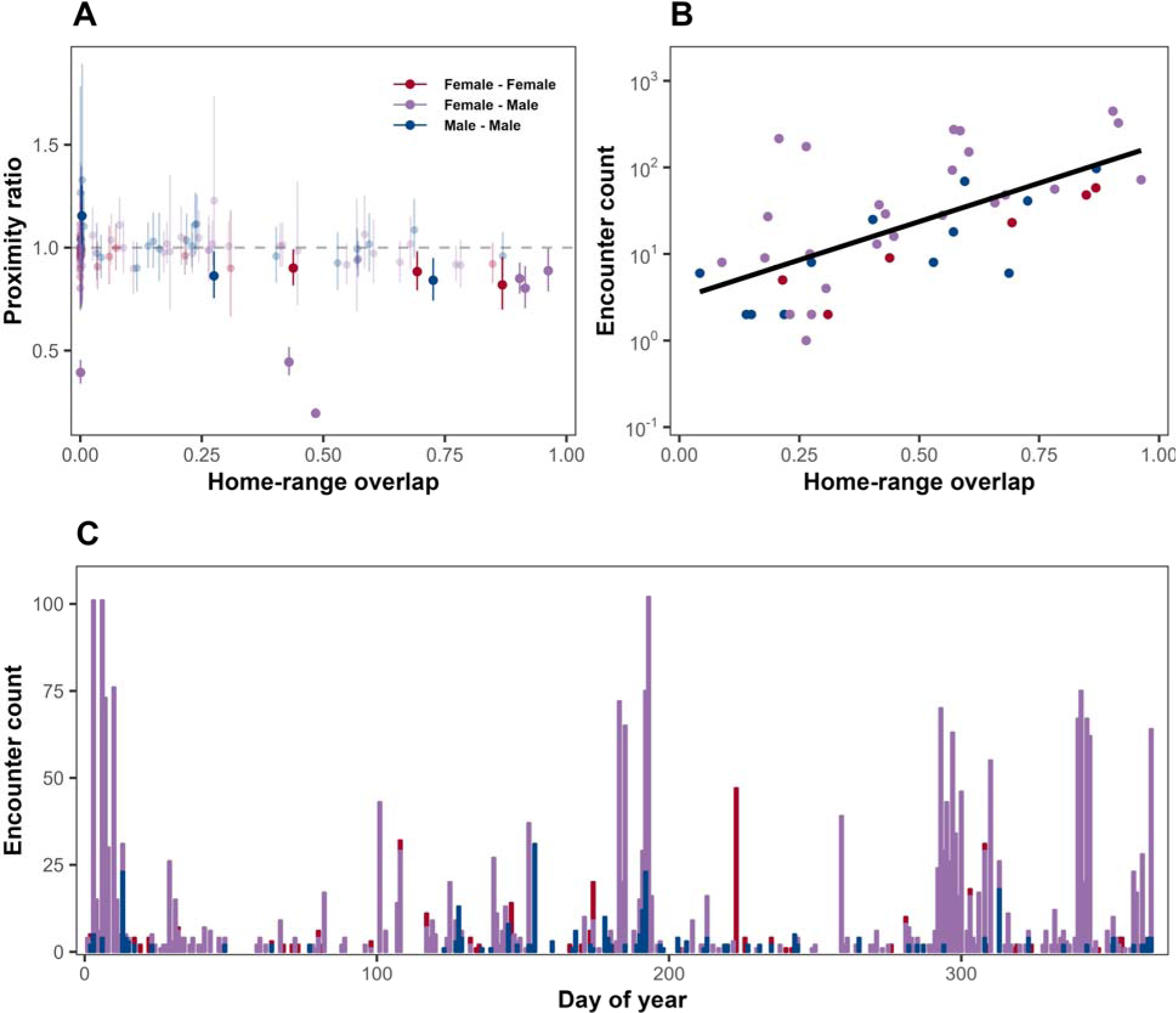
Patterns in giant anteater encounters. Panel A) shows the estimated proximity ratio values as a function of home-range overlap for the 121 dyads. A proximity ratio above 1 indicates individuals in a dyad were further apart or a proximity ratio below 1 indicates individuals in a dyad were closer on average than expected for independent movement. The 12 dyads with proximity ratios that differed significantly from the expectations under independent movement proximity ratio value are highlighted with deeper saturation of colour, the remaining are with proximity ratios of 1 are faded in colour. Panel B) depicts the total number of observed encounter events as a function of home-range overlap for the 43 dyads where at least one encounter was observed. Note that the y-axis in B) is log_10_-scaled. Panel C) shows the distribution of these encounter events over the annual cycle. Note how there is a pronounced peak in female-male encounters between ca. May and June, followed by a second peak between ca. October and January.

We identified a total of 2774 encounter events, which occurred between only 43 of the 121 dyads (284 for male-male, 145 for female-female, and 2345 for female-male). As expected, there was a positive correlation between the amount of home-range overlap and the number of observed encounters (β = 3.47 ± 0.09 SE, p < 0.01, Fig. 3B). In addition, sex had a significant effect on encounter rate, with both female-male and male-male dyads having significantly more encounters than female-female dyads (β = 1.73 ± 0.09 SE, p < 0.01; and β = 0.59 ± 0.10 SE, p < 0.01, respectively). Notably, 1079, or 38.8%, of the identified encounters were observed between the 11 dyads with proximity ratios below 1. Interestingly, we also found that while male-male and female-female encounters occurred at a relatively consistent rate throughout the year, female-male encounters had two pronounced peaks (Fig. 3C). The first occurred between ca. May and June, whereas the second occurred between ca. October and January.

### Correlated movement between giant anteaters

In general, we found that giant anteaters exhibited little to no correlation in their movement. For the 11 dyads with proximity ratios below 1, the mean amount of total correlation in their movement was 0.01 (0.00 – 0.03), with the majority of this being attributed to correlation in the diffusion component of their movement.

As a case example for these general trends, we present a more in-depth exploration for “Thomas” and “Margaret”, a female-male dyad that exhibited substantial home-range overlap (0.92; 95% CI: 0.88 – 0.94; Fig. 4A), as well as a proximity ratio which suggested significant attraction (0.80; 95% CI: 0.71 – 0.91). The mean distance between this dyad was 0.65 km (range: 0.00 – 2.06 km; Fig. 4B), and though they freely occupy the same space with each other, and encountered each other 326 times, their movement was generally independent. The mean total correlation in their movement was 0.04 (95% CI: −0.02 – 0.06), with no periods of pronounced correlation. In particular, we note a period in mid to late October where this dyad remained physically close to one another, and encountered each other 306 times over 10 days, yet still moved independently. While we only presented results for a single dyad, similar patterns were observed across all dyads (see supplementary material).

**Figure 4.**
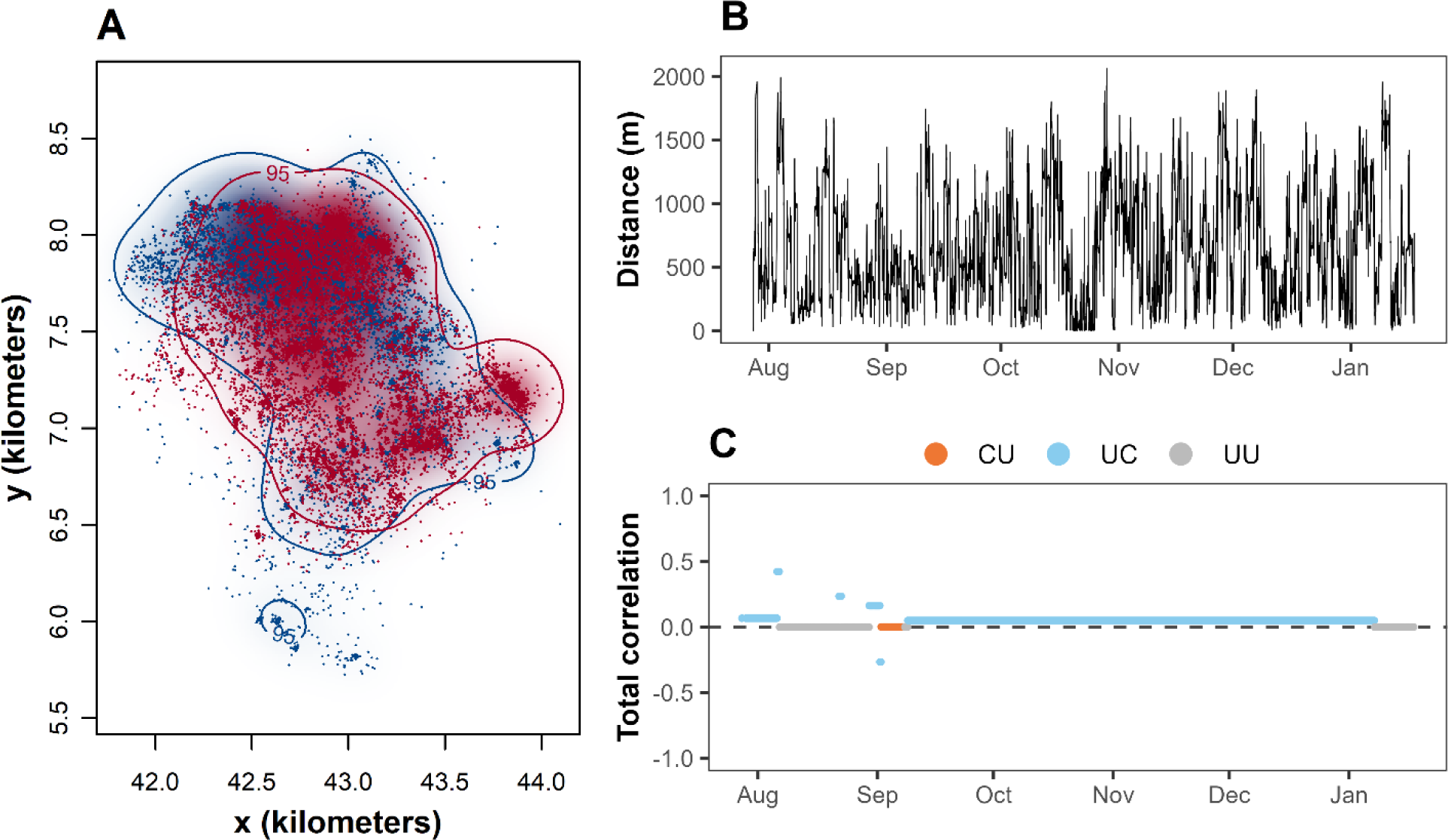
Case example of fine-scale interactions between a female-male dyad. Panel A) shows the GPS fixes with the estimated home-range probability density function and 95% contours. Blue indicates male, “Thomas” and red indicates female, “Margaret”. Panel B) depicts the Euclidean distance between the female-male between individuals over time. Panel C) shows the total correlated movement between individuals where orange (CU) indicates correlated & drift uncorrelated diffusion, light blue (UC) indicates uncorrelated & drift correlated diffusion, and grey (UU) indicates uncorrelated & drift uncorrelated diffusion. Note how there is a pronounced period of clustering where their distance is low in mid to late October in Panel B) and yet there is no correlation in their movement in Panel C) as the estimated correlation is centered around the dotted line at 0.

## Discussion

Quantifying fine-scale interactions between individuals revealed a complementary, but more nuanced picture of giant anteater social behaviour than was previously known (Shaw *et al*., 1987; Bertassoni & Milléo Costa, 2010; Júnior & Bertassoni, 2014; Catapani *et al*., 2020). While associations between dyads were generally weak, with limited evidence of attraction, avoidance, nor correlated movement, there were two seasonal peaks in female-male encounters. Furthermore, encounters between females occurred less frequently than male-male and female-male encounters. Collectively, these findings suggest giant anteaters are a largely solitary species that willingly share space with conspecifics, and that peaks in encounters are likely related to mating behaviour, though with little to no spatial segregation. Such information provides valuable insight into heretofore unknown aspects of the ecology of this iconic, but understudied species, and will help support conservation efforts to maintain viable giant anteater populations, such as reintroductions or area-based management.

### Broad-scale patterns in giant anteater social-spatial ecology

At a broad spatio-temporal scale, we found substantial overlap between individual home-ranges, with no clear structure in their spatial arrangement. Moreover, as every individual had overlap with at least one other animal, giant anteaters are unlikely to be a territorial species (Isbell *et al*., 2021; Schlichting *et al*., 2022). This suggests that either their home ranges are indefensible (Isbell *et al*., 2021), or there is no net benefit to defending the resources within their home ranges (*sensu* Grant, 1993). In this regard, it should be noted that due to the anti-predator defence mechanisms of their ant and termite prey (Lubin & Montgomery, 1981; Lubin, 1983; Redford, 1985; Naples, 1999), giant anteaters rarely, if ever, deplete a mound or nest (henceforth referred to as a “patch”) (Redford, 1985). When they are no longer able to tolerate their preys’ defenses such as biting or chemical secretions, they move on to another patch before the one they are presently feeding on has been depleted (Lubin & Montgomery, 1981; Redford, 1985). In addition to this patch switching behaviour, giant anteaters have low metabolic rates thus require lower levels of food intake than other, comparably sized mammals (McNab, 1984). When these traits are considered alongside the high abundance of ants and termites in the study region, giant anteaters likely experience high food security. The substantial overlap we observed between giant anteaters is therefore consistent with the expectation of the Resource Dispersion Hypothesis (RDH), which predicts that if resources are not limited, multiple individuals can share the same space with conspecifics (Macdonald, 1983; Macdonald & Johnson, 2015). The high amount of spatial overlap among conspecifics, alongside the low variation in home-range size, also suggests that giant anteaters are unlikely to exhibit social dominance in their habitat selection (i.e., the Ideal Despotic Distribution; Kacelnik, Krebs & Bernstein, 1992).

The observed spatial structure of giant anteater home ranges can also provide insight into the species’ mating system. In this regard, we found that the arrangement of giant anteater home ranges showed no evidence of mate guarding behaviour (i.e., male home ranges exclusively encompassing the ranges of one or more females, Clutton-Brock, 1989; Lukas & Clutton-Brock, 2012). Moreover, lack of spatial organization suggests that giant anteaters may not exhibit strong territoriality tendencies to establish home ranges for guarding mates which is indicative of female defense theory (Ostfeld, 1987; Palomares *et al*., 2017). Despite cases of marking behaviour, which can be indicative of territoriality, having been reported in giant anteaters, it may be used as other forms of communication (Braga, Santos & Batista, 2010) or it may be associated with grooming such as maintaining sharp claws to open ant and termite nests (Bertassoni, *pers. obs.*). Notably, the Brazilian Cerrado is a relatively stable environment (Werneck *et al*., 2012), and giant anteater breeding is asynchronous and can occur anytime throughout the year (Júnior & Bertassoni, 2014). Though we did notice two seasonal peaks in female-male encounters, the size and location of giant anteater home ranges remained stable throughout the year, further indicating that intrasexual competition is unlikely to be a major factor underlying the spatial structure of giant anteater home ranges. Based on kinship theory, the spatial distribution of solitary species may also be driven by relatedness, such that neighbouring conspecifics may be more tolerant of one another, willing to share space and resources, when they are closely related (Hamilton, 1964; Rogers, 1987; Minta, 1993; Støen *et al*., 2005; Smith, 2014; Elbroch *et al*., 2017). Unfortunately, however, the dispersal rates and the rate of genetic mixing within the population is currently unknown. Without insight into the genetic structure of the population, it is thus uncertain as to what extent kinship plays a role in their socio-spatial ecology and future work in this area is clearly needed. Nonetheless, when the above-discussed facts are viewed alongside our field observations, and combined with the extensive parental care that female giant anteaters reportedly provide their offspring (Eisenberg & Redford, 1999, p. 92-93; Bertassoni, *pers. obs*), we suggest that giant anteaters are likely a polygamous species (Crook, Ellis & Goss-Custard, 1976; Ostfeld, 1987; Clutton-Brock, 1989) and consistent with other findings (Desbiez *et al*., 2020).

### Fine-scale interactions between giant anteaters

Existing knowledge on giant anteater sociality is based on brief reports and are primarily from observations on captive animals (del Valle Jerez & Halloy, 2003; Bertassoni & Milléo Costa, 2010; Eyer & Miller, 2020), with only limited data on the behaviour of free-ranging animals (Montgomery & Lubin, 1975; Shaw *et al*., 1987; Bertassoni & Milléo Costa, 2010; Bertassoni *et al*., 2021; Giroux *et al*., 2021). From these, giant anteaters are described as mostly solitary in the wild except during the mating season where courtship behaviour has been observed (Shaw *et al*., 1987; Braga *et al*., 2010; Bertassoni & Desbiez, 2022). Our findings on the finer-scale interactions between individuals support these earlier works. The pairwise separation distances, proximity ratios, encounter rates, and the weak correlations between giant anteater movement revealed that individuals did not actively avoid each other, yet neither did they exhibit any cohesive movements or collective behaviour indicative of social behaviour (Giardina, 2008). For instance, we found evidence that female-male encounters occurred more regularly than both male-male and female-female encounters. Furthermore, while male-male and female-female encounters tended to occur at a fairly consistent rate throughout the year, there were two seasonal peaks in female-male encounters between May to June, and October to January. This suggests that mating is likely an important driver of giant anteater encounter rates. Notably though, associations between dyads were generally weak, with limited evidence of correlated movement. When individuals were close enough to each other such that interactions could plausibly occur, they still exhibit independent movement, as our case example highlights. The lack of any detectable avoidance behaviour that is indicative of territoriality (Isbell *et al*., 2021) or following behaviour that is indicative of mate guarding (Sherman, 1989), further support our overarching finding that giant anteaters are largely asocial (Sandell, 1989; Yoerg, 1999), and easily share space with conspecifics.

Though these findings are broadly in line with existing studies, there have been reports of courtship and following behaviour during mating season (Shaw *et al*., 1987; Júnior & Bertassoni, 2014). Furthermore, agonistic behaviour and social interactions between individuals during the mating season have also been documented (Shaw *et al*., 1987; Júnior & Bertassoni, 2014). Due to 20-minute time intervals of our GPS recordings, not all behaviour changes or events may be recorded. For example, a conflict event occurring between a pair of free-ranging females lasting approximately 4 minutes has been observed (Bertassoni, *pers. obs.*) meanwhile Kreutz, Fischer & Linsenmair (2009) observed an agonistic event lasting approximately 20 minutes between those of unknown sex. In addition, anecdotal evidence suggests that giant anteater populations have female-biased dispersal rates (Desbiez, *pers. obs.*), which is in line with the lower levels of overlap and encounters we observed between females. This suggests that there may be some amount of female-female reproductive competition (Perrin & Mazalov, 2000). Despite the current accuracy of GPS tracking data, there are still limitations and constraints, and these data alone do not always paint a complete picture of animal behaviour. Therefore, combining GPS tracking data with observational studies is crucial in further understanding the socio-spatial ecology of giant anteaters in the wild.

### Conservation implications

Giant anteaters have been experiencing population declines throughout their geographic range and are currently classified as vulnerable to extinction (Miranda *et al*., 2014). Yet, much of the species’ ecology remains unknown and these knowledge gaps have been impeding conservation efforts. Giant anteaters are experiencing extreme Human Induced Rapid Environmental Change (HIREC; Sih, 2013). In southern Brazil, they are distributed throughout cattle ranching pastures, where fire burning of the pastures multiple times annually in different areas is common practice to stimulate growth of new grasses for cattle grazing (Eloy *et al*., 2019). The practice of burning of pastures can pose serious implications for giant anteaters, especially due to their slow-moving nature and for females with a new pup. However, with the agricultural expansion of soybean crops in the Cerrado (Soterroni *et al*., 2019), many of these cattle ranching pastures are being converted into soybean crops (Cohn *et al*., 2016; Nepstad *et al*., 2019), which poses further serious implications for giant anteaters. Conventional agricultural practices involve the use of insecticides and pesticides, which not only pose a risk to giant anteaters’ health (e.g., Medici *et al*., 2021), but will almost certainly have a detrimental impact on the availability of their insect prey and thermal refuges. Consequently, the joint pressures of being condensed into smaller areas and decreased food availability may see behavioural changes radiating throughout population, despite their present capacity to share space. Moreover, changes in their social environment such as territorial due to competition and aggression or agonistic behaviour may arise due to conflict over access to resources if ant and termite prey become limiting. If giant anteater populations continue to decrease, there is potential risk for the population to become isolated and inbreeding or a genetic bottleneck may arise which can lead to decrease in genetic diversity (Collevatti *et al*., 2007; Clozato *et al*., 2017; Barragán-Ruiz *et al*., 2021). It should be noted that giant anteaters occupy an extensive biogeographic range (Miranda *et al*., 2014), where Shaw *et al*. (1987) highlight that activity patterns differ geographically. It is thus possible that our findings may be only generalizable to giant anteaters occupying ecosystems similar to the Cerrado biome. Additionally, not all individuals within the study area may have been captured and tracked, therefore, the social behaviour exhibited in this study may not reflect the whole population or species (i.e., differ across the range) To date, 43 individuals are the largest number of giant anteaters ever monitored. Subsequently, future work should aim to assess the extent to which these results are representative of giant anteater socio-spatial ecology in other parts of their range.

## Conclusion

Mammalian social systems are often based on reproductive strategies (Clutton-Brock, 2009), however, they are not wholly derived from mating behaviour (Prox & Farine, 2019). The spatial environment and movements may also be shaped or influenced by other mechanistic drivers, such as competition for resources (Montgomery & Lubin, 1975), disease dynamics (Albery *et al*., 2020), or density dependence (Smith *et al*., 2023). Using fine-scaled GPS tracking data allowed us to make inferences on the socio-spatial ecology of giant anteaters. From our analyses, we found no evidence of territoriality, nor mate guarding behaviour, suggesting that giant anteaters readily share space with conspecifics, and likely have a polygamous mating system. Such information is crucial for area-based management strategies for sustainable development while maintaining ecological integrity and the conservation of giant anteaters and their habitats. Future studies should incorporate the genetics of giant anteaters to provide further insight into socio-spatial behaviour as a consequence of kinship.

## Acknowledgements

This work was supported by the NSERC Discovery Grant RGPIN-2021-02758 to MJN as well as the Canadian Foundation for Innovation. We would like to thank the donors to the Anteaters and Highways Project especially the Foundation Segre as well as North American and European Zoos listed at http://www.giantanteater.org/. We would also like to thank the owners of all the ranches that allowed us to monitor animals on their property, in particular Nhuveira, Quatro Irmãos and Santa Lourdes ranches, and also thank D. Yogui, M. Alves, and D. Kluyber. We would like to thank all the volunteers who helped us in carrying out the fieldwork.

## Author Contributions

All authors conceived the ideas and designed methodology; ALJD collected the data; AC and MJN analysed the data; AC and MJN led the writing of the manuscript. All authors contributed critically to the drafts and gave final approval for publication.

## References

Albery, G.F., Newman, C., Ross, J.B., MacDonald, D.W., Bansal, S. & Buesching, C. (2020). Negative density-dependent parasitism in a group-living carnivore. Proc. R. Soc. B. 287, 20202655.

Allen, A.M. & Singh, N.J. (2016). Linking Movement Ecology with Wildlife Management and Conservation. Front. Ecol. Evol. 3.

Ascensão, F. & Desbiez, A.L.J. (2022). Assessing the impact of roadkill on the persistence of wildlife populations: A case study on the giant anteater. Perspectives in Ecology and Conservation 20, 272–278.

Barragán-Ruiz, C.E., Silva-Santos, R., Saranholi, B.H., Desbiez, A.L.J. & Galetti, P.M. (2021). Moderate Genetic Diversity and Demographic Reduction in the Threatened Giant Anteater, Myrmecophaga tridactyla. Front. Genet. 12, 669350.

Bartumeus, F., Da Luz, M.G.E., Viswanathan, G.M. & Catalan, J. (2005). Animal search strategies: a quantitative random-walk analysis. Ecology 86, 3078–3087.

Bates, D., Mächler, M., Bolker, B. & Walker, S. (2015). Fitting Linear Mixed-Effects Models Using **lme4**. J. Stat. Soft. 67.

Bertassoni, A., Bianchi, R.D.C. & Desbiez, A.L.J. (2021). Giant Anteater Population Density Estimation and Viability Analysis Through MotionLSensitive Camera Records. Jour. Wild. Mgmt. 85, 1554–1562.

Bertassoni, A. & Desbiez, A.L.J. (2022). The Imperiled Giant Anteater: Ecology and Conservation. In Imperiled: The Encyclopedia of Conservation: 166–176. Elsevier.

Bertassoni, A. & Milléo Costa, L.C. (2010). Behavioral repertoire of giant anteater (Myrmecophaga tridactyla, Linnaeus 1758) in nature at Serra da Canastra National Park, MG and in captivity at Curitiba Zoo. Rev. etol. 9.

Bertassoni, A., Mourão, G. & Bianchi, R.D.C. (2020). Space use by giant anteaters (*Myrmecophaga tridactyla*) in a protected area within humanLmodified landscape. Ecol Evol 10, 7981–7994.

Bertassoni, A. & Ribeiro, M.C. (2019). Space use by the giant anteater (Myrmecophaga tridactyla): a review and key directions for future research. Eur J Wildl Res 65, 93.

Braga, F.G., Santos, R.E.F. & Batista, A.C. (2010). Marking behavior of the giant anteater Myrmecophaga tridactyla (Mammalia: Myrmecophagidae) in Southern Brazil. Zoologia (Curitiba, Impr.) 27, 07–12.

Bro-Jørgensen, J., Franks, D.W. & Meise, K. (2019). Linking behaviour to dynamics of populations and communities: application of novel approaches in behavioural ecology to conservation. Phil. Trans. R. Soc. B 374, 20190008.

Brooks, M., E., Kristensen, K., Benthem, K., J., van, Magnusson, A., Berg, C., W., Nielsen, A., Skaug, H., J., Mächler, M. & Bolker, B., M. (2017). glmmTMB Balances Speed and Flexibility Among Packages for Zero-inflated Generalized Linear Mixed Modeling. The R Journal 9, 378.

Burt, W.H. (1943). Territoriality and Home Range Concepts as Applied to Mammals. Journal of Mammalogy 24, 346.

Calabrese, J.M. & Fleming, C.H. (2023). corrMove: Analyze Correlated Movements in Multi-Individual Datasets.

Calabrese, J.M., Fleming, C.H., Fagan, W.F., Rimmler, M., Kaczensky, P., Bewick, S., Leimgruber, P. & Mueller, T. (2018). Disentangling social interactions and environmental drivers in multi-individual wildlife tracking data. Phil. Trans. R. Soc. B 373, 20170007.

Calabrese, J.M., Fleming, C.H. & Gurarie, E. (2016). ctmm: an R package for analyzing animal relocation data as a continuousLtime stochastic process. Methods Ecol Evol 7, 1124–1132.

Catapani, M., Theodoro Molina, K., Martins Costa Lopes, A. & Miranda, F. (2020). Report of three non-agonistic encounters of free-living giant anteaters (Myrmecophaga tridactyla). Edentata: The Newsletter of the IUCN/SSC Anteater, Sloth and Armadillo Specialist Group 31–34.

Clozato, C.L., Miranda, F.R., Lara-Ruiz, P., Collevatti, R.G. & Santos, F.R. (2017). Population structure and genetic diversity of the giant anteater (Myrmecophaga tridactyla: Myrmecophagidae, Pilosa) in Brazil. Genet. Mol. Biol. 40, 50–60.

Clutton-Brock, T. (2009). Structure and function in mammalian societies. Phil. Trans. R. Soc. B 364, 3229–3242.

Clutton-Brock, T.H. (1989). Review Lecture: Mammalian mating systems. Proceedings of the Royal Society of London. Series B, Biological sciences 236.

Cohn, A.S., Gil, J., Berger, T., Pellegrina, H. & Toledo, C. (2016). Patterns and processes of pasture to crop conversion in Brazil: Evidence from Mato Grosso State. Land Use Policy 55, 108–120.

Collevatti, R.G., Leite, K.C.E., Miranda, G.H.B.D. & Rodrigues, F.H.G. (2007). Evidence of high inbreeding in a population of the endangered giant anteater, Myrmecophaga tridactyla (Myrmecophagidae), from Emas National Park, Brazil. Genet. Mol. Biol. 30, 112–120.

Crook, J.H., Ellis, J.E. & Goss-Custard, J.D. (1976). Mammalian social systems: Structure and function. Animal Behaviour 24, 261–274.

Desbiez, A.L.J., Bertassoni, A. & Traylor-Holzer, K. (2020). Population viability analysis as a tool for giant anteater conservation. Perspectives in Ecology and Conservation 18, 124–131.

Di Blanco, Y.E., Jiménez Pérez, I. & Di Bitetti, M.S. (2015). Habitat selection in reintroduced giant anteaters: the critical role of conservation areas. JMAMMAL 96, 1024–1035.

Dougherty, E.R., Seidel, D.P., Carlson, C.J., Spiegel, O. & Getz, W.M. (2018). Going through the motions: incorporating movement analyses into disease research. Ecol Lett 21, 588–604.

Eisenberg, J.F. & Redford, K.H. (1999). The Central Neotropics: Ecuador, Peru, Bolivia, Brazil, Mammals of the Neotropics. 1st edn. Chicago: University of Chicago Press.

Elbroch, L.M., Levy, M., Lubell, M., Quigley, H. & Caragiulo, A. (2017). Adaptive social strategies in a solitary carnivore. Sci. Adv. 3, e1701218.

Eloy, L., Schmidt, I.B., Borges, S.L., Ferreira, M.C. & Dos Santos, T.A. (2019). Seasonal fire management by traditional cattle ranchers prevents the spread of wildfire in the Brazilian Cerrado. Ambio 48, 890–899.

Eyer, A.E. & Miller, L.J. (2020). Evaluating the Influence of Conspecifics on a Male Giant Anteater’s (Myrmecophaga tridactyla) Pacing Behavior. AB&C 7, 556–566.

Fleming, C.H. & Calabrese, J.M. (2017). A new kernel density estimator for accurate homeLrange and speciesLrange area estimation. Methods Ecol Evol 8, 571–579.

Fleming, C.H. & Calabrese, J.M. (2022). ctmm: Continuous-Time Movement Modeling.

Fleming, C.H., Fagan, W.F., Mueller, T., Olson, K.A., Leimgruber, P. & Calabrese, J.M. (2015). Rigorous home range estimation with movement data: a new autocorrelated kernel density estimator. Ecology 96, 1182–1188.

Fraser, K.C., Davies, K.T.A., Davy, C.M., Ford, A.T., Flockhart, D.T.T. & Martins, E.G. (2018). Tracking the Conservation Promise of Movement Ecology. Front. Ecol. Evol. 6, 150.

Giardina, I. (2008). Collective behavior in animal groups: Theoretical models and empirical studies. HFSP Journal 2, 205–219.

Giroux, A., Ortega, Z., Oliveira-Santos, L.G.R., Attias, N., Bertassoni, A. & Desbiez, A.L.J. (2021). Sexual, allometric and forest cover effects on giant anteaters’ movement ecology. PLoS ONE 16, e0253345.

Grant, J.W.A. (1993). Whether or not to defend? The influence of resource distribution. Marine Behaviour and Physiology 23, 137–153.

Hamilton, W.D. (1964). The genetical evolution of social behaviour. I. Journal of Theoretical Biology 7, 1–16.

Hertel, A.G., Niemelä, P.T., Dingemanse, N.J. & Mueller, T. (2020). A guide for studying among-individual behavioral variation from movement data in the wild. Movement Ecology 8, 30.

Isbell, L.A., Bidner, L.R., Loftus, J.C., Kimuyu, D.M. & Young, T.P. (2021). Absentee owners and overlapping home ranges in a territorial species. Behav Ecol Sociobiol 75, 21.

Jesmer, B.R., Merkle, J.A., Goheen, J.R., Aikens, E.O., Beck, J.L., Courtemanch, A.B., Hurley, M.A., McWhirter, D.E., Miyasaki, H.M., Monteith, K.L. & Kauffman, Matthew. J. (2018). Is ungulate migration culturally transmitted? Evidence of social learning from translocated animals. Science 361, 1023–1025.

Júnior, J.F.M. & Bertassoni, A. (2014). Potential Agonistic Courtship and Mating Behavior between Two Adult Giant Anteaters (*Myrmecophaga tridactyla*). Edentata 15, 69–72.

Kacelnik, A., Krebs, J.R. & Bernstein, C. (1992). The ideal free distribution and predator-prey populations. Trends in Ecology & Evolution 7, 50–55.

Kays, R., Crofoot, M.C., Jetz, W. & Wikelski, M. (2015). Terrestrial animal tracking as an eye on life and planet. Science 348, aaa2478.

Kluyber, D., Attias, N., Alves, M.H., Alves, A.C., Massocato, G. & Desbiez, A.L.J. (2021). Physical capture and chemical immobilization procedures for a mammal with singular anatomy: the giant anteater (Myrmecophaga tridactyla). Eur J Wildl Res 67, 67.

Kreutz, K., Fischer, F. & Linsenmair, K.E. (2009). Observations of lntraspecific Aggression in Giant Anteaters (*Myrmecophaga tridactyla*). Edentata 8–10, 6–7.

Leblond, M., Dussault, C. & Ouellet, J.-P. (2013). Impacts of Human Disturbance on Large Prey Species: Do Behavioral Reactions Translate to Fitness Consequences? PLoS ONE 8, e73695.

Lubin, Y.D. (1983). Eating ants is no picnic. Nature History 92, 54–59.

Lubin, Y.D. & Montgomery, G.G. (1981). Defenses of Nasutitermes Termites (Isoptera, Termitidae) Against Tamandua Anteaters (Edentata, Myrmecophagidae). Biotropica 13, 66.

Lukas, D. & Clutton-Brock, T. (2012). Life histories and the evolution of cooperative breeding in mammals. Proc. R. Soc. B. 279, 4065–4070.

Macdonald, D.W. (1983). The ecology of carnivore social behaviour. Nature 301, 379–384.

Macdonald, D.W. & Johnson, D.D.P. (2015). Patchwork planet: the resource dispersion hypothesis, society, and the ecology of life. J Zool 295, 75–107.

Martinez-Garcia, R., Fleming, C.H., Seppelt, R., Fagan, W.F. & Calabrese, J.M. (2020). How range residency and long-range perception change encounter rates. Journal of Theoretical Biology 498, 110267.

McNab, B.K. (1984). Physiological convergence amongst antLeating and termiteLeating mammals. Journal of Zoology 203, 485–510.

Medici, E.P., Fernandes-Santos, R.C., Testa-José, C., Godinho, A.F. & Brand, A.-F. (2021). Lowland tapir exposure to pesticides and metals in the Brazilian Cerrado. Wildl. Res. 48, 393–403.

Minta, S.C. (1993). Sexual differences in spatio-temporal interaction among badgers. Oecologia 96, 402–409.

Miranda, F., Bertassoni, A. & Abba, A.M. (2014). Myrmecophaga tridactyla (No. e. T14224A47441961)., The IUCN Red List of Threatened Species. International Union for Conservation of Nature and Natural Resources.

Montgomery, G.G. & Lubin, Y.D. (1975). Prey influences on movements of neotropical anteaters. In : 103–131. Phillips, R.L. & Jonkel, C.J. (Eds.). Presented at the Proceedings of the 1975 Predator Symposium, Montana Forest and Conservation Experiment Station, University of Montana, Missoula.

Moorcroft, P.R., Lewis, M.A. & Crabtree, R.L. (1999). Home Range Analysis Using a Mechanistic Home Range Model. Ecology 80, 1656–1665.

Mourão, G. & Medri, Í.M. (2007). Activity of a specialized insectivorous mammal (Myrmecophaga tridactyla) in the Pantanal of Brazil. Journal of Zoology 271, 187–192.

Mueller, T., O’Hara, R.B., Converse, S.J., Urbanek, R.P. & Fagan, W.F. (2013). Social Learning of Migratory Performance. Science 341, 999–1002.

Naples, V.L. (1999). Morphology, evolution and function of feeding in the giant anteater (*Myrmecophaga tridactyla*). Journal of Zoology 249, 19–41.

Nathan, R., Getz, W.M., Revilla, E., Holyoak, M., Kadmon, R., Saltz, D. & Smouse, P.E. (2008). A movement ecology paradigm for unifying organismal movement research. Proc. Natl. Acad. Sci. U.S.A. 105, 19052–19059.

Nathan, R., Monk, C.T., Arlinghaus, R., Adam, T., Alós, J., Assaf, M., Baktoft, H., Beardsworth, C.E., Bertram, M.G., Bijleveld, A.I., Brodin, T., Brooks, J.L., Campos-Candela, A., Cooke, S.J., Gjelland, K.Ø., Gupte, P.R., Harel, R., Hellström, G., Jeltsch, F., Killen, S.S., Klefoth, T., Langrock, R., Lennox, R.J., Lourie, E., Madden, J.R., Orchan, Y., Pauwels, I.S., Říha, M., Roeleke, M., Schlägel, U.E., Shohami, D., Signer, J., Toledo, S., Vilk, O., Westrelin, S., Whiteside, M.A. & Jarić, I. (2022). Big-data approaches lead to an increased understanding of the ecology of animal movement. Science 375, eabg1780.

Nepstad, L.S., Gerber, J.S., Hill, J.D., Dias, L.C.P., Costa, M.H. & West, P.C. (2019). Pathways for recent Cerrado soybean expansion: extending the soy moratorium and implementing integrated crop livestock systems with soybeans. Environ. Res. Lett. 14, 044029.

Noonan, M.J., Ascensão, F., Yogui, D.R. & Desbiez, A.L.J. (2022). Roads as ecological traps for giant anteaters. Animal Conservation 25, 182–194.

Noonan, M.J., MartinezLGarcia, R., Davis, G.H., Crofoot, M.C., Kays, R., Hirsch, B.T., Caillaud, D., Payne, E., Sih, A., Sinn, D.L., Spiegel, O., Fagan, W.F., Fleming, C.H. & Calabrese, J.M. (2021). Estimating encounter location distributions from animal tracking data. Methods Ecol Evol 12, 1158–1173.

Noonan, M.J., Martinez-Garcia, R., Fleming, C.H., De Figueiredo, B.G., Ali, A.H., Attias, N., Belant, J.L., Beyer, D.E., Berteaux, D., Bidner, L.R., Boone, R., Boutin, S., Brito, J., Brown, M., Carter, A., Castellanos, A., Castellanos, F.X., Chitwood, C., Darlington, S., De La Torre, J.A., Dekker, J., DePerno, C., Droghini, A., Farhadinia, M., Fennessy, J., Fichtel, C., Ford, A., Gill, R., Goheen, J.R., Oliveira-Santos, L.G.R., Hebblewhite, M., Hodges, K.E., Isbell, L.A., Janssen, R., Kappeler, P., Kays, R., Kaczensky, P., Kauffman, M., LaPoint, S., Lashley, M.A., Leimgruber, P., Little, A., Macdonald, D.W., Masiaine, S., McBride, R.T., Medici, E.P., Mertes, K., Moorman, C., Morato, R.G., Mourão, G., Mueller, T., Neilson, E.W., Pastorini, J., Patterson, B.D., Pereira, J., Petroelje, T.R., Piecora, K., Power, R.J., Rachlow, J., Ranglack, D.H., Roshier, D., Safford, K., Scott, D.M., Serrouya, R., Songer, M., Songsasen, N., Stabach, J., Stacy-Dawes, J., Swingen, M.B., Thompson, J., Tucker, M.A., Velilla, M., Yarnell, R.W., Young, J., Fagan, W.F. & Calabrese, J.M. (2023). The search behavior of terrestrial mammals (preprint). Ecology.

Noonan, M.J., Tucker, M.A., Fleming, C.H., Akre, T.S., Alberts, S.C., Ali, A.H., Altmann, J., Antunes, P.C., Belant, J.L., Beyer, D., Blaum, N., BöhningLGaese, K., Cullen, L., De Paula, R.C., Dekker, J., DrescherLLehman, J., Farwig, N., Fichtel, C., Fischer, C., Ford, A.T., Goheen, J.R., Janssen, R., Jeltsch, F., Kauffman, M., Kappeler, P.M., Koch, F., LaPoint, S., Markham, A.C., Medici, E.P., Morato, R.G., Nathan, R., OliveiraLSantos, L.G.R., Olson, K.A., Patterson, B.D., Paviolo, A., Ramalho, E.E., Rösner, S., Schabo, D.G., Selva, N., Sergiel, A., Xavier Da Silva, M., Spiegel, O., Thompson, P., Ullmann, W., Zięba, F., ZwijaczLKozica, T., Fagan, W.F., Mueller, T. & Calabrese, J. M. (2019). A comprehensive analysis of autocorrelation and bias in home range estimation. Ecological Monographs 89.

Ostfeld, R.S. (1987). On the Distinction between Female Defense and Resource Defense Polygyny. Oikos 48, 238.

Palomares, F., Lucena-Pérez, M., López-Bao, J.V. & Godoy, J.A. (2017). Territoriality ensures paternity in a solitary carnivore mammal. Sci Rep 7, 4494.

Perrin, N. & Mazalov, V. (2000). Local Competition, Inbreeding, and the Evolution of SexLBiased Dispersal. The American Naturalist 155, 116–127.

Powell, R.A. (2000). Animal home ranges and territories and home range estimators. In Research techniques in animal ecology: controversies and consequences, Methods and cases in conservation science: 65–110. Boitani, L. & Fuller, T.K. (Eds.). New York: Columbia University Press.

Prox, L. & Farine, D. (2019). A framework for conceptualizing dimensions of social organization in mammals. Ecol Evol 10, 791–807.

R Core Team. (2022). R: A language and environment for statistical computing.

Redford, K.H. (1985). Feeding and food preference in captive and wild Giant anteaters *(Myrmecophaga tridactyla)*. Journal of Zoology 205, 559–572.

Rogers, L.L. (1987). Effects of Food Supply and Kinship on Social Behavior, Movements, and Population Growth of Black Bears in Northeastern Minnesota. Wildlife Monographs 3–72.

Sandell, M. (1989). The Mating Tactics and Spacing Patterns of Solitary Carnivores. In Carnivore Behavior, Ecology, and Evolution: 164–182. Gittleman, J.L. (Ed.). Boston, MA: Springer US.

Schick, R.S., Loarie, S.R., Colchero, F., Best, B.D., Boustany, A., Conde, D.A., Halpin, P.N., Joppa, L.N., McClellan, C.M. & Clark, J.S. (2008). Understanding movement data and movement processes: current and emerging directions. Ecology Letters 11, 1338–1350.

Schlichting, P.E., Boughton, R.K., Anderson, W., Wight, B., VerCauteren, K.C., Miller, R.S. & Lewis, J.S. (2022). Seasonal variation in space use and territoriality in a large mammal (Sus scrofa). Sci Rep 12, 4023.

Shaw, J.H., Machado-Neto, J. & Carter, T.S. (1987). Behavior of Free-Living Giant Anteaters (Myrmecophaga tridactyla). Biotropica 19, 255.

Sherman, P.W. (1989). Mate guarding as paternity insurance in Idaho ground squirrels. Nature 338, 418–420.

Shiratsuru, S., Studd, E.K., Boutin, S., Peers, M.J.L., Majchrzak, Y.N., Menzies, A.K., Derbyshire, R., Jung, T.S., Krebs, C.J., Boonstra, R. & Murray, D.L. (2023). When death comes: linking predator–prey activity patterns to timing of mortality to understand predation risk. Proc. R. Soc. B. 290, 20230661.

Sih, A. (2013). Understanding variation in behavioural responses to human-induced rapid environmental change: a conceptual overview. Animal Behaviour 85, 1077–1088.

Smith, B.J., MacNulty, D.R., Stahler, D.R., Smith, D.W. & Avgar, T. (2023). DensityLdependent habitat selection alters drivers of population distribution in northern Yellowstone elk. Ecology Letters 26, 245–256.

Smith, J.E. (2014). Hamilton’s legacy: kinship, cooperation and social tolerance in mammalian groups. Animal Behaviour 92, 291–304.

Smithson, M. & Verkuilen, J. (2006). A better lemon squeezer? Maximum-likelihood regression with beta-distributed dependent variables. Psychological Methods 11, 54–71.

Soterroni, A.C., Ramos, F.M., Mosnier, A., Fargione, J., Andrade, P.R., Baumgarten, L., Pirker, J., Obersteiner, M., Kraxner, F., Câmara, G., Carvalho, A.X.Y. & Polasky, S. (2019). Expanding the Soy Moratorium to Brazil’s Cerrado. Sci. Adv. 5, eaav7336.

Souza, C.M., Z. Shimbo, J., Rosa, M.R., Parente, L.L., A. Alencar, A., Rudorff, B.F.T., Hasenack, H., Matsumoto, M., G. Ferreira, L., Souza-Filho, P.W.M., De Oliveira, S.W., Rocha, W.F., Fonseca, A.V., Marques, C.B., Diniz, C.G., Costa, D., Monteiro, D., Rosa, E.R., Vélez-Martin, E., Weber, E.J., Lenti, F.E.B., Paternost, F.F., Pareyn, F.G.C., Siqueira, J.V., Viera, J.L., Neto, L.C.F., Saraiva, M.M., Sales, M.H., Salgado, M.P.G., Vasconcelos, R., Galano, S., Mesquita, V.V. & Azevedo, T. (2020). Reconstructing Three Decades of Land Use and Land Cover Changes in Brazilian Biomes with Landsat Archive and Earth Engine. Remote Sensing 12, 2735.

Støen, O.-G., Bellemain, E., Sæbø, S. & Swenson, J.E. (2005). Kin-related spatial structure in brown bears Ursus arctos. Behav Ecol Sociobiol 59, 191–197.

Strandburg-Peshkin, A., Farine, D.R., Couzin, I.D. & Crofoot, M.C. (2015). Shared decision-making drives collective movement in wild baboons. Science 348, 1358–1361.

Strassburg, B.B.N., Brooks, T., Feltran-Barbieri, R., Iribarrem, A., Crouzeilles, R., Loyola, R., Latawiec, A.E., Oliveira Filho, F.J.B., Scaramuzza, C.A.D.M., Scarano, F.R., Soares-Filho, B. & Balmford, A. (2017). Moment of truth for the Cerrado hotspot. Nat Ecol Evol 1, 0099.

Tórrez-Herrera, L.L., Davis, G.H. & Crofoot, M.C. (2020). Do Monkeys Avoid Areas of Home Range Overlap Because They Are Dangerous? A Test of the Risk Hypothesis in White-Faced Capuchin Monkeys (Cebus capucinus). Int J Primatol 41, 246–264.

del Valle Jerez, S. & Halloy, M. (2003). El oso hormiguero, Myrmecophaga tridactyla: crecimiento e independización de una cría. Mastozoología Neotropical 10, 323–330.

Webber, Q.M.R., Albery, G.F., Farine, D.R., PinterLWollman, N., Sharma, N., Spiegel, O., Vander Wal, E. & Manlove, K. (2023). Behavioural ecology at the spatial–social interface. Biological Reviews brv.12934.

Werneck, F.P., Nogueira, C., Colli, G.R., Sites, J.W. & Costa, G.C. (2012). Climatic stability in the Brazilian Cerrado: implications for biogeographical connections of South American savannas, species richness and conservation in a biodiversity hotspot: Climatic stability and biodiversity in the Cerrado. Journal of Biogeography 39, 1695–1706.

Whittington, J., Hebblewhite, M., Baron, R.W., Ford, A.T. & Paczkowski, J. (2022). Towns and trails drive carnivore movement behaviour, resource selection, and connectivity. Mov Ecol 10, 17.

Wickham, H. (2016). ggplot2: Elegant Graphics for Data Analysis. Springer-Verlag New York.

Winner, K., Noonan, M.J., Fleming, C.H., Olson, K.A., Mueller, T., Sheldon, D. & Calabrese, J.M. (2018). Statistical inference for home range overlap. Methods Ecol Evol 9, 1679–1691.

Yoerg, S.I. (1999). Solitary is not Asocial: Effects of Social Contact in Kangaroo Rats (Heteromyidae: Dipodomys heermanni). Ethology 105, 317–333.

