## Appendix S1 for "The socio-spatial ecology of giant anteaters in the Brazilian Cerrado": Anteater_Appendix.html

###### This document was created on October 02, 2023.

- **Overview**
- **Data**
- **Movement
  Models**
- **Home Range Overlap**
  - Estimating home range
    areas
  - Home range
    results
  - Estimating home range
    overlap
  - Home range overlap results
- **Proximity and
  Encounters**
  - Proximity
    - Estimating proximity ratio
    - Proximity ratio results
  - Encounters
    - Estimating distance
    - Sensitivity analysis
    - Estimating encounters
    - Encounter
      results
- **Correlated Movement**
  - Exploring deviations
    in proximity ratios
  - Deviated pairs results
  - Estimating distances
    in the deviated pairs
  - Estimating correlated
    movement
  - Correlated movement
    results
  - Case study
- **Figures**
  - Figure 1
  - Figure 2
  - Figure 3
  - Figure 4
- End Session
- References

---

### **Overview**

Our primary aim was to understand the movement ecology of giant
anteaters (*Myrmecophaga tridactyla*) at the social-spatial
interface. Specifically, we aimed to address two overarching
questions:

- Is the spatial arrangement of giant anteater home ranges governed by
  social factors (e.g., territoriality, mating groups, social groups,
  etc.)?
- Does giant anteater movement within their home range exhibit signs
  of social interactions (e.g., avoidance, attraction, correlated
  movement, etc.)?

In order to answer our two core research questions, we carried out
three separate analyses focused on estimating patterns in:

1. home-range overlap;
2. proximity and encounter rates; and
3. correlated movement.

To do this, we used GPS data to quantify key socio-spatial patterns
using the methods implemented in the `ctmm` and the
`corrMove` packages in `R`. In this appendix, we
describe the workflow and specific steps that was used to carry out in
each of these analyses below.

Data visualisation and analysis were carried out using R (version
4.2.2, R Core Team, 2022) and the following R packages: ctmm (version
1.1.0, Fleming & Calabrese, 2022), lme4 (version 1.1.31, Bates et
al., 2015), glmmTMB (version 1.1.7, Brooks et al., 2017), corrMove
(version 0.1.0, Calabrese & Fleming, 2023), and ggplot2 (version
3.4.2, Wickham, 2016). All R scripts can be found in the GitHub
repository at https://github.com/QuantitativeEcologyLab/Giant\_Anteater.

---

### **Data**

Before analysis, we calibrated the GPS measurement error and removed
any outlying data points (for full details on data pre-processing see
Appendix S2 in (Noonan et al., 2022). First step of the analysis is to
import the data and convert to a telemetry object for use in the
`ctmm` package.

```
#load packages
library(readr)
library(ggplot2)
library(dplyr)           
library(tidyr)    
library(tibble)
library(lubridate)       
library(gridExtra)       
library(scales)          
library(osmdata)
```

```
## Warning: package 'osmdata' was built under R version 4.3.1
```

```
library(terra)           
library(tidyterra)
```

```
## Warning: package 'tidyterra' was built under R version 4.3.1
```

```
library(sf)              
library(ctmm)            
library(lme4)
library(glmmTMB)
library(corrMove)
```

```
#import GPS data
DATA_GPS <- read_csv("data/Anteaters_NoOutliers.csv")

#import supplementary data containing biological information
DATA_META <- read_csv("data/Anteater_Results_Final.csv")

#correct mismatch ID entries
DATA_GPS$ID[DATA_GPS$ID == "Larry 267"] <- "Larry"
DATA_GPS$ID[DATA_GPS$ID == "Little Rick"] <- "Little_Rick"

#correct mismatch ID entries
DATA_META$ID[DATA_META$ID == "Little Rick"] <- "Little_Rick"
```

While a total of 43 individuals were collared as part of the larger
monitoring effort (Noonan et al., 2022), here we restricted our analyses
to 23 range-resident individuals living in three separate clusters.
These individuals were selected as they resided in areas where there was
high confidence that all resident giant anteaters were equipped with GPS
trackers. The resulting dataset consisted of 528,324 GPS fixes.

```
#subset to range-resident individuals
GPS_df <- DATA_GPS[which(DATA_GPS$ID %in% 
                           c("Alexander", "Annie", "Anthony", "Beto", "Bumpus",
                             "Cate", "Christoffer","Elaine", "Hannah","Jackson",
                             "Jane","Kyle", "Larry", "Little_Rick", "Luigi",
                             "Makao", "Margaret", "Maria", "Puji", "Reid", 
                             "Rodolfo", "Sheron", "Thomas")),]

#subset to range-resident individuals
bio_df <- DATA_META[c(1:3,8:10,12,14,17,19,20,22,23,25:29,33:35,37,38),]
#subset the biological data
bio_df <- bio_df[,c(1:3,5)]

#add site location to the dataframe
bio_df$Site <- NA
bio_df$Site[bio_df$Road == "MS-040"] <- 1
bio_df$Site[bio_df$Road == "BR_267"] <- 2
bio_df
```

```
## # A tibble: 23 × 5
##    ID          Sex    Age      Road    Site
##    <chr>       <chr>  <chr>    <chr>  <dbl>
##  1 Alexander   Male   Adult    MS-040     1
##  2 Annie       Female Adult    BR_267     2
##  3 Anthony     Male   Subadult MS-040     1
##  4 Beto        Male   Adult    BR_267     2
##  5 Bumpus      Female Adult    MS-040     1
##  6 Cate        Female Adult    MS-040     1
##  7 Christoffer Male   Adult    MS-040     1
##  8 Elaine      Female Adult    MS-040     1
##  9 Hannah      Female Adult    BR_267     2
## 10 Jackson     Male   Adult    MS-040     1
## # ℹ 13 more rows
```

```
#convert GPS dataset to a telemetry object
DATA_TELEMETRY <- as.telemetry(GPS_df)

#summary of the dataset
summary(DATA_TELEMETRY)
```

```
##             interval (min) period (mon) longitude  latitude
## Alexander               20    11.921256 -53.75980 -21.14109
## Annie                   20    11.096325 -53.49576 -21.63346
## Anthony                 20    14.080832 -53.75685 -21.11174
## Beto                    20    12.045447 -53.50511 -21.63142
## Bumpus                  20    13.232697 -53.75040 -21.14285
## Cate                    20    12.362887 -53.75986 -21.12229
## Christoffer             20     9.007152 -53.75909 -21.10599
## Elaine                  20    13.136653 -53.75061 -21.11452
## Hannah                  20    11.245092 -53.59645 -21.66039
## Jackson                 20    14.463984 -53.74615 -21.15241
## Jane                    20    11.162642 -53.48273 -21.62628
## Kyle                    20    14.387329 -53.75067 -21.14322
## Larry                   20     6.830496 -53.49571 -21.63331
## Little_Rick             20    10.232655 -53.75634 -21.15988
## Luigi                   20    11.888605 -53.58243 -21.64823
## Makao                   20    12.184317 -53.75183 -21.14762
## Margaret                20     8.828889 -53.50742 -21.62868
## Maria                   20    14.664479 -53.60082 -21.63678
## Puji                    20    12.045415 -53.75366 -21.14839
## Reid                    20    11.674806 -53.50453 -21.63001
## Rodolfo                 20    13.414102 -53.74969 -21.13223
## Sheron                  20     6.436336 -53.60102 -21.63810
## Thomas                  20     5.887029 -53.50656 -21.62690
```

```
#visualisation of the data
plot(DATA_TELEMETRY)
```

---

### **Movement Models**

After preparing the data, we fit the continuous-time movement models
following the workflow described in Calabrese et al. (2016).

```
#fit movement models
GUESS <- lapply(DATA_TELEMETRY[1:23], function(b) ctmm.guess(b,interactive=FALSE))
FIT <- lapply(1:23, function(i) ctmm.select(DATA_TELEMETRY[[i]],GUESS[[i]]))
names(FIT) <- names(DATA_TELEMETRY[1:23])
overlap(FIT)
```

```
#summary of the fitted model
summary(FIT)
```

```
##                ΔAICc ΔRMSPE (m) DOF[area]
## Thomas           0.0  113.85920 269.95105
## Sheron       14373.6  228.02211 146.83169
## Larry        27151.7  176.65376 236.89821
## Christoffer 165078.4  490.26622 214.64362
## Jane        210689.0   93.98707 328.07112
## Reid        215804.9    0.00000 301.48633
## Annie       215859.4  118.87981 463.29147
## Little_Rick 224929.3  725.14962 163.38726
## Hannah      226745.9  394.07773 261.28251
## Beto        233868.1  164.37743 312.60590
## Luigi       259010.9 1941.61466  47.45471
## Alexander   278336.4  149.77752 312.63782
## Puji        285987.2   82.08636 429.92689
## Makao       291710.2  202.29586 302.00316
## Cate        302045.1  146.34625 456.13202
## Maria       323204.1   19.68532 320.84660
## Bumpus      341042.9  127.04284 386.26756
## Anthony     357897.3   72.62748 379.72961
## Jackson     377618.2  295.29036 364.69938
## Rodolfo     384332.7  607.04544 309.30108
## Kyle        431957.5  324.42868 561.17180
## Margaret    100601.1  103.65326 231.11540
## Elaine      368562.0  698.47717  93.38052
```

---

### **Home Range Overlap**

#### Estimating home range areas

We estimated giant anteater home ranges using Autocorrelated Kernel
Density Estimation (AKDE; Fleming et al., 2015) using the
`ctmm` package. AKDE, corrects for autocorrelation induced
bias (Noonan et al., 2019) by conditioning the bandwidth optimisation on
the data’s autocorrelation structure (Fleming & Calabrese, 2017).
Thus, home range estimation first required fitting a series of
continuous-time movement models to the GPS tracking data and identifying
the best model via small sample-sized corrected Akaike’s Information
Criterion (AICc) (Fleming & Calabrese, 2022). Giant anteater home
ranges were then estimated conditional on each individual’s best-fit
movement model.

```
#calculate AKDE home range estimates based on the best fit model, create aligned UDs
AKDE <- akde(DATA_TELEMETRY[1:23],FIT)
overlap(AKDE)
```

#### Home range results

```
#create a dataframe to store home range area statistics from the AKDE
HR_size <- data.frame()

#loop through each object in the AKDE list
for (i in 1:length(AKDE)) {
  #extract the home range area statistics from summary
  summary <- summary(AKDE[[i]])$CI
  
  #bind the summary to the dataframe
  HR_size <- rbind(HR_size, as.data.frame(summary))
}

row.names(HR_size) <- NULL

#add biological data to dataframe
HR_size <- cbind(HR_size, bio_df)
HR_size <- relocate(HR_size, c(low, est, high), .after = Site)
names(HR_size)[6] <- "HR_low"
names(HR_size)[7] <- "HR_est"
names(HR_size)[8] <- "HR_high"
```

```
#calculate the mean total home range size
round(mean(HR_size$HR_est), 2)
```

```
## [1] 5.45
```

```
#calculate CIs of the mean total home range size
round(mean(HR_size$HR_low), 2)
```

```
## [1] 4.55
```

```
round(mean(HR_size$HR_high), 2)
```

```
## [1] 6.45
```

Here, we calculate the home range statistics based on a sex parameter
using the `ctmm` function `meta()`.

```
#does home range size differ between sexes?

#subset each individual based on their sex
AKDE_male <- AKDE[c("Alexander", "Anthony", "Beto","Christoffer","Jackson",
                    "Kyle", "Larry", "Little_Rick", "Luigi", "Reid", 
                    "Rodolfo", "Thomas")]
AKDE_female <- AKDE[c("Annie", "Bumpus", "Cate", "Elaine", "Hannah",
                      "Jane","Makao", "Margaret", "Maria", "Puji",
                      "Sheron")]

#calculate mean home range size for male
meta(AKDE_male)
```

```
##                     ΔAICc
## inverse-Gaussian    0.000
## Dirac-δ          1608.435
```

```
##                   low      est      high
## mean (km²)  3.6943465 7.277281 12.937360
## CoV² (RVAR) 0.3006536 1.242143  2.838906
## CoV  (RSTD) 0.5675121 1.153527  1.743883
```

```
#calculate mean home range size for female
meta(AKDE_female)
```

```
##                     ΔAICc
## inverse-Gaussian   0.0000
## Dirac-δ          288.9002
```

```
##                    low       est      high
## mean (km²)  2.56403589 3.3229394 4.2340351
## CoV² (RVAR) 0.05286221 0.1735889 0.3647909
## CoV  (RSTD) 0.23610028 0.4278434 0.6202201
```

```
#test to see significance of sex on home range
AKDE_sex_compare <- list(male = AKDE_male,
                         female = AKDE_female)
COL_sex <- c("#004488", "#A50026")
meta(AKDE_sex_compare, col = COL_sex, sort = TRUE)
```

```
## * Sub-population male
```

```
##                     ΔAICc
## inverse-Gaussian    0.000
## Dirac-δ          1608.435
```

```
## * Sub-population female
```

```
##                     ΔAICc
## inverse-Gaussian   0.0000
## Dirac-δ          288.9002
```

```
## * Joint population
```

```
##                     ΔAICc
## inverse-Gaussian    0.000
## Dirac-δ          2264.043
```

```
## * Joint population versus sub-populations (best models)
```

```
##                     ΔAICc
## Sub-population   0.000000
## Joint population 3.587004
```

```
## , , low
## 
##             /male  /female
## male/   1.0000000 0.950149
## female/ 0.1868822 1.000000
## 
## , , est
## 
##             /male  /female
## male/   1.0000000 2.153857
## female/ 0.4073595 1.000000
## 
## , , high
## 
##             /male /female
## male/   1.0000000 3.93176
## female/ 0.8460534 1.00000
```

---

#### Estimating home range overlap

Home range overlap was estimated for all pairs of individuals (i.e.,
dyads) via the Bhattacharyya coefficient (Winner et al., 2018).

```
#subset home range overlap based on the site location
AKDE_1 <- AKDE[c("Alexander", "Anthony", "Bumpus", "Cate", "Christoffer",
                 "Elaine", "Jackson", "Kyle", "Little_Rick", "Makao",
                 "Puji", "Rodolfo")]

AKDE_2 <- AKDE[c("Annie", "Beto", "Hannah", "Jane", "Larry",
                 "Luigi", "Margaret", "Maria", "Reid", "Sheron",
                 "Thomas")]
```

```
#calculate 95% AKDE home range overlap for a pairwise comparison for each site
overlap_1 <- overlap(AKDE_1, level = 0.95)
overlap_2 <- overlap(AKDE_2, level = 0.95)
```

```
#create a pairwise dataframe by pairing up every individual at each site

#extract CI 'est' matrix from array
overlap_1_est <- overlap_1$CI[ , , 2]
#remove duplicate values of the matrix
overlap_1_est[upper.tri(overlap_1_est, diag = TRUE)] <- NA
#Create a new data frame based on the overlap values
overlap_1_df <- as.data.frame(overlap_1_est)
overlap_1_df$anteater_A <- rownames(overlap_1_df)
overlap_1_df <- pivot_longer(overlap_1_df, cols = -anteater_A, 
                             names_to = 'anteater_B', values_to = 'overlap_est', values_drop_na = TRUE)

#extract CI 'low' matrix from array
overlap_1_low <- overlap_1$CI[ , , 1]
#remove duplicate values of the matrix
overlap_1_low[upper.tri(overlap_1_low, diag = TRUE)] <- NA
#create a new dataframe based on the overlap values
overlap_1_low <- as.data.frame(overlap_1_low)
overlap_1_low$anteater_A <- rownames(overlap_1_low)
overlap_1_low <- pivot_longer(overlap_1_low, cols = -anteater_A, 
                              names_to = 'anteater_B', values_to = 'overlap_low',
                              values_drop_na = TRUE)
overlap_1_df <- left_join(overlap_1_df, overlap_1_low, 
                          by = c("anteater_A", "anteater_B"))
overlap_1_df <- relocate(overlap_1_df, overlap_low, .before = overlap_est)

#extract CI 'high' matrix from array
overlap_1_high <- overlap_1$CI[ , , 3]
#remove duplicate values of the matrix
overlap_1_high[upper.tri(overlap_1_high, diag = TRUE)] <- NA
#create a new dataframe based on the overlap values
overlap_1_high <- as.data.frame(overlap_1_high)
overlap_1_high$anteater_A <- rownames(overlap_1_high)
overlap_1_high <- pivot_longer(overlap_1_high, cols = -anteater_A, 
                               names_to = 'anteater_B', values_to = 'overlap_high',
                               values_drop_na = TRUE)
overlap_1_df <- left_join(overlap_1_df, overlap_1_high, 
                          by = c("anteater_A", "anteater_B"))

#add biological data to dataframe
overlap_1_df <- left_join(overlap_1_df, rename(bio_df, anteater_A = ID), 
                          by = "anteater_A")
overlap_1_df <- left_join(overlap_1_df, rename(bio_df, anteater_B = ID), 
                          by = "anteater_B", suffix = c(".A", ".B"))
#add column to indicate which sexes that are being compared
overlap_1_df <- mutate(overlap_1_df,
                       sex_comparison = 
                         case_when(paste(Sex.A, Sex.B) == "Male Male" ~ "male-male",
                                   paste(Sex.A, Sex.B) == "Female Female" ~ "female-female",
                                   paste(Sex.A, Sex.B) == "Male Female" ~ "female-male",
                                   paste(Sex.A, Sex.B) == "Female Male" ~ "female-male"))
#assign site
overlap_1_df$site <- 1
overlap_1_df <- relocate(overlap_1_df, 
                         c("overlap_low", "overlap_est", "overlap_high"), .after = site)
```

```
#extract CI 'est' matrix from array
overlap_2_est <- overlap_2$CI[ , , 2]
#remove duplicate values of the matrix
overlap_2_est[upper.tri(overlap_2_est, diag = TRUE)] <- NA
#create a new data frame based on the overlap values
overlap_2_df <- as.data.frame(overlap_2_est)
overlap_2_df$anteater_A <- rownames(overlap_2_df)
overlap_2_df <- pivot_longer(overlap_2_df, cols = -anteater_A, 
                             names_to = 'anteater_B', values_to = 'overlap_est', values_drop_na = TRUE)

#extract CI 'low' matrix from array
overlap_2_low <- overlap_2$CI[ , , 1]
#remove duplicate values of the matrix
overlap_2_low[upper.tri(overlap_2_low, diag = TRUE)] <- NA
#create a new data frame based on the overlap values
overlap_2_low <- as.data.frame(overlap_2_low)
overlap_2_low$anteater_A <- rownames(overlap_2_low)
overlap_2_low <- pivot_longer(overlap_2_low, cols = -anteater_A, 
                              names_to = 'anteater_B', values_to = 'overlap_low',
                              values_drop_na = TRUE)
overlap_2_df <- left_join(overlap_2_df, overlap_2_low, 
                          by = c("anteater_A", "anteater_B"))
overlap_2_df <- relocate(overlap_2_df, overlap_low, .before = overlap_est)

#extract CI 'high' matrix from array
overlap_2_high <- overlap_2$CI[ , , 3]
#remove duplicate values of the matrix
overlap_2_high[upper.tri(overlap_2_high, diag = TRUE)] <- NA
#create a new data frame based on the overlap values
overlap_2_high <- as.data.frame(overlap_2_high)
overlap_2_high$anteater_A <- rownames(overlap_2_high)
overlap_2_high <- pivot_longer(overlap_2_high, cols = -anteater_A, 
                               names_to = 'anteater_B', values_to = 'overlap_high',
                               values_drop_na = TRUE)
overlap_2_df <- left_join(overlap_2_df, overlap_2_high, 
                          by = c("anteater_A", "anteater_B"))

#add biological data to dataframe
overlap_2_df <- left_join(overlap_2_df, rename(bio_df, anteater_A = ID), 
                          by = "anteater_A")
overlap_2_df <- left_join(overlap_2_df, rename(bio_df, anteater_B = ID), 
                          by = "anteater_B", suffix = c(".A", ".B"))
#add column to indicate which sexes that are being compared
overlap_2_df <- mutate(overlap_2_df,
                       sex_comparison = 
                         case_when(paste(Sex.A, Sex.B) == "Male Male" ~ "male-male",
                                   paste(Sex.A, Sex.B) == "Female Female" ~ "female-female",
                                   paste(Sex.A, Sex.B) == "Male Female" ~ "female-male",
                                   paste(Sex.A, Sex.B) == "Female Male" ~ "female-male"))
#assign site
overlap_2_df$site <- 2
overlap_2_df <- relocate(overlap_2_df, 
                         c("overlap_low", "overlap_est", "overlap_high"), .after = site)
```

```
#combine both sites into one dataframe
overlap_df <- rbind(overlap_1_df, overlap_2_df)
overlap_df$pair_ID <- paste(overlap_df$anteater_A, overlap_df$anteater_B, sep = "_")
overlap_df <- relocate(overlap_df, pair_ID, .before = anteater_A)

#clean up environment
rm(overlap_1_low, overlap_1_est, overlap_1_high,
   overlap_2_low, overlap_2_est, overlap_2_high)
```

#### Home range overlap results

```
#calculate mean total home range overlap 
round(mean(overlap_df$overlap_est), 2)
```

```
## [1] 0.2
```

```
#calculate range of total home range overlap 
round(min(overlap_df$overlap_est), 2)
```

```
## [1] 0
```

```
round(max(overlap_df$overlap_est), 2)
```

```
## [1] 0.96
```

To determine if sex was a factor underpinning the degree of pairwise
overlap, we fit a Generalized Linear Mixed Model (GLMM) with a beta
distribution and a logit link function to the home range overlap
estimates, with pairwise sex as a predictor variable (i.e., male-male,
female-female, and female-male). In addition, site was included as a
random effect. Because the overlap values ranged between [0,1], we
rescaled the values as (y(n-1) + 0.5)/n in order for them to lie within
the (0,1) interval (Smithson & Verkuilen, 2006). This model was then
compared to a similar model that excluded the pairwise sex predictor
variable using a likelihood ratio test.

```
#rescale the values
min_val <- min(overlap_df$overlap_est)
max_val <- max(overlap_df$overlap_est)
squeeze_min <- 0.001
squeeze_max <- 0.999
overlap_df$overlap_est_squeezed <- ((overlap_df$overlap_est - min_val) / (max_val - min_val)) * (squeeze_max - squeeze_min) + squeeze_min
overlap_df <- relocate(overlap_df, overlap_est_squeezed, .after = overlap_high)

#test for significance in sex, compare model with and without sex as a variable
HRO_test <- glmmTMB(overlap_est_squeezed ~ sex_comparison + (1|site), 
                    family = beta_family(link = "logit"), data = overlap_df)
HRO_test2 <- glmmTMB(overlap_est_squeezed ~ 1 + (1|site), 
                     family = beta_family(link = "logit"), data = overlap_df)
HRO_test_results <- anova(HRO_test, HRO_test2)
HRO_test_pvalue <- round(HRO_test_results$`Pr(>Chisq)`[2], 2)
```

Home range overlap results based on sex comparisons (i.e.. male-male,
female-female, and female-male).

```
#number of home range overlap in each sex comparison category
table(overlap_df$sex_comparison)
```

```
## 
## female-female   female-male     male-male 
##            25            65            31
```

```
#calculate mean home range overlap and the range based on sex comparisons
round(mean(overlap_df$overlap_est[overlap_df$sex_comparison == "male-male"]), 2)
```

```
## [1] 0.22
```

```
round(min(overlap_df$overlap_est[overlap_df$sex_comparison == "male-male"]), 2)
```

```
## [1] 0
```

```
round(max(overlap_df$overlap_est[overlap_df$sex_comparison == "male-male"]), 2)
```

```
## [1] 0.87
```

```
round(mean(overlap_df$overlap_est[overlap_df$sex_comparison == "female-female"]), 2)
```

```
## [1] 0.14
```

```
round(min(overlap_df$overlap_est[overlap_df$sex_comparison == "female-female"]), 2)
```

```
## [1] 0
```

```
round(max(overlap_df$overlap_est[overlap_df$sex_comparison == "female-female"]), 2)
```

```
## [1] 0.87
```

```
round(mean(overlap_df$overlap_est[overlap_df$sex_comparison == "female-male"]), 2)
```

```
## [1] 0.21
```

```
round(min(overlap_df$overlap_est[overlap_df$sex_comparison == "female-male"]), 2)
```

```
## [1] 0
```

```
round(max(overlap_df$overlap_est[overlap_df$sex_comparison == "female-male"]), 2)
```

```
## [1] 0.96
```

---

### **Proximity and Encounters**

While home-range overlap describes patterns in spatial structuring,
it does not directly indicate whether individuals are likely to be in
the same place at the same time (Winner et al., 2018). In order to
understand the temporal component of giant anteaters’ socio-spatial
behaviour, we estimated a proximity ratio for all dyads via the
`ctmm` function `proximity()`.

#### Proximity

##### Estimating proximity ratio

The proximity ratio was estimated by comparing a dyad’s observed
separation distances with the separation distances expected under
completely random movement. A proximity ratio of 1 is thus consistent
with independent movement, values <1 indicate that the two
individuals are closer on average than expected for independent
movement, and vice versa for values >1.

```
#subset movement models of individuals based on their site location
FIT_1 <- FIT[c("Alexander", "Anthony", "Bumpus", "Cate", "Christoffer",
               "Elaine", "Jackson", "Kyle", "Little_Rick", "Makao",
               "Puji", "Rodolfo")]
FIT_2 <- FIT[c("Annie", "Beto", "Hannah", "Jane", "Larry",
               "Luigi", "Margaret", "Maria", "Reid", "Sheron",
               "Thomas")]
```

```
#create empty columns for the results to be saved to
overlap_1_df$proximity_low <- NA
overlap_1_df$proximity_est <- NA
overlap_1_df$proximity_high <- NA

#calculate the proximity statistics
for(i in 1:nrow(overlap_1)){
  ANIMAL_A <- as.character(overlap_1_df[i, 'anteater_A'])
  ANIMAL_B <- as.character(overlap_1_df[i, 'anteater_B'])
  
  TRACKING_DATA.1 <- DATA_TELEMETRY[c(ANIMAL_A, ANIMAL_B)]
  MODELS.1 <- list(FIT_1[ANIMAL_A][[1]], FIT_1[ANIMAL_B][[1]])
  
  PROXIMITY1 <- tryCatch(
    {
      #calculate the proximity statistic
      PROXIMITY_1 <- proximity(data = TRACKING_DATA.1, CTMM = MODELS.1, GUESS=ctmm(error=FALSE))},
    error=function(err){
      PROXIMITY_1 <- c(NA,NA,NA)
      return(PROXIMITY_1)
    }
  )
  
  overlap_1_df[i, c("proximity_low")] <- PROXIMITY_1[1]
  overlap_1_df[i, c("proximity_est")] <- PROXIMITY_1[2]
  overlap_1_df[i, c("proximity_high")] <- PROXIMITY_1[3]
  
  #save results to a csv file
  write.csv(overlap_1_df, "data/DATA_proximity_1.csv", row.names = FALSE)
  #track progress
  cat("finished index", i, "\n")
}
```

```
#create empty columns for the results to be saved to
overlap_2_df$proximity_low <- NA
overlap_2_df$proximity_est <- NA
overlap_2_df$proximity_high <- NA

#calculate the proximity statistics
for(i in 1:nrow(overlap_2_df)){
  ANIMAL_A <- as.character(overlap_2_df[i, 'anteater_A'])
  ANIMAL_B <- as.character(overlap_2_df[i, 'anteater_B'])
  
  TRACKING_DATA_2 <- DATA_TELEMETRY[c(ANIMAL_A, ANIMAL_B)] 
  MODELS_2 <- list(FIT_2[ANIMAL_A][[1]], FIT_2[ANIMAL_B][[1]])
  
  PROXIMITY_2 <- tryCatch(
    {
      #calculate the proximity statistic
      PROXIMITY_2 <- proximity(data = TRACKING_DATA_2, 
                               CTMM = MODELS_2, 
                               GUESS=ctmm(error=FALSE))},
    error=function(err){
      PROXIMITY_2 <- c(NA,NA,NA)
      return(PROXIMITY_2)
    }
  )
  overlap_2_df[i, c("proximity_low")] <- PROXIMITY_2[1]
  overlap_2_df[i, c("proximity_est")] <- PROXIMITY_2[2]
  overlap_2_df[i, c("proximity_high")] <- PROXIMITY_2[3]
  
  #save results in a csv file
  write.csv(overlap_2_df, "data/DATA_proximity_2.csv", row.names = FALSE)
  #track progress
  cat("finished index", i, "\n")
}
```

```
#import proximity statistics data
DATA_proximity_1 <- read_csv("data/DATA_proximity_1.csv")
DATA_proximity_2 <- read_csv("data/DATA_proximity_2.csv")

#correct mismatch entry
DATA_proximity_1$anteater_A[DATA_proximity_1$anteater_A == "Little Rick"] <- "Little_Rick"
DATA_proximity_1$anteater_B[DATA_proximity_1$anteater_B == "Little Rick"] <- "Little_Rick"
DATA_proximity_2$anteater_A[DATA_proximity_2$anteater_A == "Larry 267"] <- "Larry"
DATA_proximity_2$anteater_B[DATA_proximity_2$anteater_B == "Larry 267"] <- "Larry"

#add missing site column to dataframe for site 2
DATA_proximity_2$site <- 2
DATA_proximity_2 <- relocate(DATA_proximity_2, site, .before = proximity_low)

#create a proximity dataframe
proximity_df <- bind_rows(DATA_proximity_1, DATA_proximity_2)
proximity_df <- proximity_df[,-3]
proximity_df <- mutate(proximity_df,
                       sex_comparison = 
                         case_when(paste(Sex.A, Sex.B) == "Male Male" ~ "male-male",
                                   paste(Sex.A, Sex.B) == "Female Female" ~ "female-female",
                                   paste(Sex.A, Sex.B) == "Male Female" ~ "female-male",
                                   paste(Sex.A, Sex.B) == "Female Male" ~ "female-male"))

#add home range overlap data to proximity dataframe
proximity_df <- left_join(overlap_df, proximity_df, by = c("anteater_A", "anteater_B",
                                                           "Sex.A", "Sex.B",
                                                           "Age.A", "Age.B",
                                                           "sex_comparison",
                                                           "site"))
```

##### Proximity ratio results

We used a GLMM with a gamma distribution, a log link function, and
site as a random effect to determine whether the estimated proximity
ratios differed between pairwise sex combinations. This model was then
compared to similar model that excluded the pairwise sex predictor
variable using a likelihood ratio test.

```
#test for significance in sex, compare model with and without sex as a variable
proximity_test <- glmer(proximity_est ~ sex_comparison + (1|site), 
                        family = Gamma(link = "log"), data = proximity_df)
```

```
## boundary (singular) fit: see help('isSingular')
```

```
proximity_test2 <- glmer(proximity_est ~ 1 + (1|site), 
                         family = Gamma(link = "log"), data = proximity_df)
```

```
## boundary (singular) fit: see help('isSingular')
```

```
proximity_test_results <- anova(proximity_test, proximity_test2)
proximity_test_pvalue <- round(proximity_test_results$`Pr(>Chisq)`[2], 2)

#test for significance in home range overlap, compare model with and without overlap as a variable
prox_overlap_test <- glmer(proximity_est ~ overlap_est + (1|site), 
                           family = Gamma(link = "log"), data = proximity_df)
```

```
## boundary (singular) fit: see help('isSingular')
```

```
prox_overlap_test2 <- glmer(proximity_est ~ 1 + (1|site), 
                            family = Gamma(link = "log"), data = proximity_df)
```

```
## boundary (singular) fit: see help('isSingular')
```

```
prox_overlap_test_results <- anova(prox_overlap_test, prox_overlap_test2)
prox_overlap_test_pvalue <- round(prox_overlap_test_results$`Pr(>Chisq)`[2], 2)
```

---

#### Encounters

In order to calculate the number of encounter events, we needed to
estimate the Euclidean distance between the individuals in each dyads at
each timepoint using the `ctmm` function
`distances()`. From these separation distances we estimated
the number of encounter events for each dyad using a 15m distance
threshold. We also performed a sensitivity analysis on the 15m
threshold.

##### Estimating distance

```
#create empty columns for the results to be saved to
proximity_df$distance_low <- NA
proximity_df$distance_est <- NA
proximity_df$distance_high <- NA
#create empty list for the results to be saved to
RES <- list()

#calculate the distance statistics
for (i in 1:nrow(overlap_df)) {
  ANIMAL_A <- as.character(overlap_df[i, 'anteater_A']) 
  ANIMAL_B <- as.character(overlap_df[i, 'anteater_B'])
  TRACKING_DATA <- DATA_TELEMETRY[c(ANIMAL_A, ANIMAL_B)]
  MODELS <- list(FIT[[ANIMAL_A]], FIT[[ANIMAL_B]])
  
  DISTANCES_RES <- tryCatch({
    distances_result <- distances(data = TRACKING_DATA, 
                                  CTMM = MODELS, 
                                  GUESS = ctmm(error = FALSE))
    data.frame(pair_ID = paste(ANIMAL_A, ANIMAL_B, sep = "_"),
               distance_low = distances_result$low, 
               distance_est = distances_result$est, 
               distance_high = distances_result$high,
               t = distances_result$t,
               timestamp = distances_result$timestamp)
  }, error = function(err) {
    data.frame(pair_ID = paste(ANIMAL_A, ANIMAL_B, sep = "_"),
               distance_low = NA,
               distance_est = NA,
               distance_high = NA,
               t = NA, 
               timestamp = NA)
  })
  
  RES[[i]] <- DISTANCES_RES
  
  #track progress
  cat("finished index", i, "\n")
}

#turn the list of list into a dataframe
DATA_DISTANCE <- do.call(rbind, RES)
```

```
#import the distance statistics data
DATA_DISTANCE <- readRDS("data/rds/DATA_DISTANCE.rds")

#locate NA values within the dataframe
DATA_DISTANCE[!complete.cases(DATA_DISTANCE), ]
#drop the 3 fixes that had no distance values 
DATA_DISTANCE <- na.omit(DATA_DISTANCE)

#add overlap and proximity information to the distance dataframe
distance_df <- merge(DATA_DISTANCE, proximity_df, by = "pair_ID")
distance_df <- relocate(distance_df, c(distance_low, distance_est, distance_high,
                                       t, timestamp), .after = proximity_high)
```

##### Sensitivity analysis

```
#set encounter radius
enc_radius <- 0:1000
enc_count <- vector("numeric", length(enc_radius))

#calculate the number of encounters occurring within each radius size
for(i in 1:length(enc_radius)){
  enc_count[i] <- sum(distance_df$distance_est < enc_radius[i])
}
```

```
plot_enc_radius_count <- 
  ggplot() +
  geom_line(aes(x = enc_radius, y = enc_count)) +
  labs(x = "Encounter radius (m)", 
       y = "Encounter count (x1000)") +
  scale_y_continuous(labels = function(enc_count) enc_count / 1000) +
  theme_bw() +
  theme(panel.grid.major = element_blank(),
        panel.grid.minor = element_blank(),
        plot.title = element_text( size = 14, family = "sans", face = "bold"),
        axis.title.y = element_text(size=10, family = "sans", face = "bold"),
        axis.title.x = element_text(size=10, family = "sans", face = "bold"),
        axis.text.y = element_text(size=8, family = "sans"),
        axis.text.x  = element_text(size=8, family = "sans"),
        legend.position="none",
        panel.background = element_rect(fill = "transparent"),
        legend.background = element_rect(fill = "transparent"),
        plot.background = element_rect(fill = "transparent", color = NA),
        plot.margin = unit(c(0.2,0.1,0.2,0.2), "cm"))
```

```
#sensitivity analysis on female-male encounter significance
#create empty columns for the results to be saved to
encounter_radius_pvalue <- vector("numeric", length(enc_radius))
pair_ID <- unique(overlap_df$pair_ID)

#loop over encounter radii
for(i in 1:length(enc_radius)){
  
  res <- list()
  
  for (j in pair_ID){
    subset_A <- distance_df[distance_df$pair_ID == j,]
    
    #count the number of times "distance_est" is below some threshold distance i 
    encounter_count <- sum(subset_A$distance_est < enc_radius[i])
    
    #save results
    res[[j]] <- data.frame(encounter_count = encounter_count,
                           overlap_est = subset_A$overlap_est[1],
                           sex_comparison = subset_A$sex_comparison[1],
                           site = subset_A$site[1])
    
  }
  
  res <- do.call(rbind, res)
  encounter_radius_test <- try(glmer(encounter_count ~ 
                                       overlap_est + sex_comparison + (1|site), 
                                     family = poisson(link = "log"), 
                                     data = res, subset = res > 0))
  encounter_radius_test2 <- try(glmer(encounter_count ~ 1 + (1|site), 
                                      family = poisson(link = "log"), 
                                      data = res, subset = res > 0))
  encounter_radius_test_results <- try(anova(encounter_radius_test, encounter_radius_test2))
  p_val <- try(encounter_radius_test_results$`Pr(>Chisq)`[2])
  encounter_radius_pvalue[i] <- ifelse(class(p_val) == "try-error", NA, p_val)
  
  #track progress
  cat("finished index", i, "\n")
}

#create an encounter radius dataframe
encounter_radius_df <- data.frame(x = enc_radius,
                                  y = encounter_radius_pvalue)

saveRDS(encounter_radius_df, file = "rds/encounter_radius_df.rds")
```

```
plot_enc_radius_pvalue <-
ggplot() +
  geom_line(data = encounter_radius_df, 
            aes(x = x, y = y),
            size = 0.15) +
  geom_hline(yintercept = 0.05, linetype = "dashed", color = "red") +
  xlab("Encounter radius (m)") +
  ylab("p-value") +
  theme_bw() +
  theme(panel.grid.major = element_blank(),
        panel.grid.minor = element_blank(),
        plot.title = element_text(size = 14, family = "sans", face = "bold"),
        axis.title.y = element_text(size=10, family = "sans", face = "bold"),
        axis.title.x = element_text(size=10, family = "sans", face = "bold"),
        axis.text.y = element_text(size=8, family = "sans"),
        axis.text.x = element_text(size=8, family = "sans"),
        panel.background = element_rect(fill = "transparent"),
        plot.background = element_rect(fill = "transparent", color = NA),
        plot.margin = unit(c(0.1,0.1,0.05,0.2), "cm")) #top, right, bot, left)
```

##### Estimating encounters

```
#create an empty column for the results to be saved to
proximity_df$encounter_count <- NA
pair_ID <- unique(proximity_df$pair_ID)

#calculate total encounters of all individuals based on sex comparison type
for (i in pair_ID){
  subset_A <- distance_df[distance_df$pair_ID == i,]
  
  #count the number of times distance is below 15
  encounter_count <- sum(subset_A$distance_est < 15)
  
  #save results
  proximity_df[proximity_df$pair_ID == i, "encounter_count"] <- encounter_count
  
}

#number of pairs that had 0 encounters
proximity_df[proximity_df$encounter_count == 0,]
```

```
## # A tibble: 78 × 21
##    pair_ID    anteater_A anteater_B Sex.A Age.A Road.A Site.A Sex.B Age.B Road.B
##    <chr>      <chr>      <chr>      <chr> <chr> <chr>   <dbl> <chr> <chr> <chr> 
##  1 Anthony_A… Anthony    Alexander  Male  Suba… MS-040      1 Male  Adult MS-040
##  2 Bumpus_An… Bumpus     Anthony    Fema… Adult MS-040      1 Male  Suba… MS-040
##  3 Cate_Alex… Cate       Alexander  Fema… Adult MS-040      1 Male  Adult MS-040
##  4 Cate_Bump… Cate       Bumpus     Fema… Adult MS-040      1 Fema… Adult MS-040
##  5 Christoff… Christoff… Alexander  Male  Adult MS-040      1 Male  Adult MS-040
##  6 Christoff… Christoff… Bumpus     Male  Adult MS-040      1 Fema… Adult MS-040
##  7 Elaine_Al… Elaine     Alexander  Fema… Adult MS-040      1 Male  Adult MS-040
##  8 Elaine_Bu… Elaine     Bumpus     Fema… Adult MS-040      1 Fema… Adult MS-040
##  9 Jackson_A… Jackson    Anthony    Male  Adult MS-040      1 Male  Suba… MS-040
## 10 Jackson_C… Jackson    Cate       Male  Adult MS-040      1 Fema… Adult MS-040
## # ℹ 68 more rows
## # ℹ 11 more variables: Site.B <dbl>, sex_comparison <chr>, site <dbl>,
## #   overlap_low <dbl>, overlap_est <dbl>, overlap_high <dbl>,
## #   overlap_est_squeezed <dbl>, proximity_low <dbl>, proximity_est <dbl>,
## #   proximity_high <dbl>, encounter_count <int>
```

```
#number of pairs that had at least 1 encounter
proximity_df[proximity_df$encounter_count != 0,]
```

```
## # A tibble: 43 × 21
##    pair_ID    anteater_A anteater_B Sex.A Age.A Road.A Site.A Sex.B Age.B Road.B
##    <chr>      <chr>      <chr>      <chr> <chr> <chr>   <dbl> <chr> <chr> <chr> 
##  1 Bumpus_Al… Bumpus     Alexander  Fema… Adult MS-040      1 Male  Adult MS-040
##  2 Cate_Anth… Cate       Anthony    Fema… Adult MS-040      1 Male  Suba… MS-040
##  3 Christoff… Christoff… Anthony    Male  Adult MS-040      1 Male  Suba… MS-040
##  4 Christoff… Christoff… Cate       Male  Adult MS-040      1 Fema… Adult MS-040
##  5 Elaine_An… Elaine     Anthony    Fema… Adult MS-040      1 Male  Suba… MS-040
##  6 Elaine_Ca… Elaine     Cate       Fema… Adult MS-040      1 Fema… Adult MS-040
##  7 Elaine_Ch… Elaine     Christoff… Fema… Adult MS-040      1 Male  Adult MS-040
##  8 Jackson_A… Jackson    Alexander  Male  Adult MS-040      1 Male  Adult MS-040
##  9 Jackson_B… Jackson    Bumpus     Male  Adult MS-040      1 Fema… Adult MS-040
## 10 Kyle_Bump… Kyle       Bumpus     Male  Suba… MS-040      1 Fema… Adult MS-040
## # ℹ 33 more rows
## # ℹ 11 more variables: Site.B <dbl>, sex_comparison <chr>, site <dbl>,
## #   overlap_low <dbl>, overlap_est <dbl>, overlap_high <dbl>,
## #   overlap_est_squeezed <dbl>, proximity_low <dbl>, proximity_est <dbl>,
## #   proximity_high <dbl>, encounter_count <int>
```

```
#calculate the number of encounters based on threshold of 15m
sum(proximity_df$encounter_count)
```

```
## [1] 2774
```

```
sum(proximity_df$encounter_count[proximity_df$sex_comparison == "male-male"])
```

```
## [1] 284
```

```
sum(proximity_df$encounter_count[proximity_df$sex_comparison == "female-female"])
```

```
## [1] 145
```

```
sum(proximity_df$encounter_count[proximity_df$sex_comparison == "female-male"])
```

```
## [1] 2345
```

##### Encounter results

We used a GLMM with a Poisson distribution, a log link function, and
site as a random effect to determine whether encounter rates differed
between the pairwise sex combinations. This model was then compared to
similar model that excluded the pairwise sex predictor variable using a
likelihood ratio test. Additionally, we used a Hierarchical Generalized
Additive Model (HGAM) with a Poisson distribution, a log link function,
to test for any temporal trends in encounters.

```
#effect of sex and overlap on encounter rates and does not include 0 encounter counts
encounter_test <- glmer(encounter_count ~ overlap_est + sex_comparison + (1|site), 
                        family = poisson(link = "log"), 
                        data = proximity_df, subset = encounter_count > 0)
encounter_test2 <- glmer(encounter_count ~ 1 + (1|site), 
                         family = poisson(link = "log"), 
                         data = proximity_df, subset = encounter_count > 0)
encounter_test_results <- anova(encounter_test, encounter_test2)
encounter_test_pvalue <- round(encounter_test_results$`Pr(>Chisq)`[2], 2)

#amount of home range overlap and the number of observed encounters
summary(encounter_test)
```

```
## Generalized linear mixed model fit by maximum likelihood (Laplace
##   Approximation) [glmerMod]
##  Family: poisson  ( log )
## Formula: encounter_count ~ overlap_est + sex_comparison + (1 | site)
##    Data: proximity_df
##  Subset: encounter_count > 0
## 
##      AIC      BIC   logLik deviance df.resid 
##   2240.5   2249.3  -1115.3   2230.5       38 
## 
## Scaled residuals: 
##     Min      1Q  Median      3Q     Max 
## -10.143  -4.244  -2.358   1.768  29.732 
## 
## Random effects:
##  Groups Name        Variance Std.Dev.
##  site   (Intercept) 0.2073   0.4553  
## Number of obs: 43, groups:  site, 2
## 
## Fixed effects:
##                           Estimate Std. Error z value Pr(>|z|)    
## (Intercept)                0.68327    0.34040   2.007   0.0447 *  
## overlap_est                3.46592    0.09053  38.283  < 2e-16 ***
## sex_comparisonfemale-male  1.73111    0.08608  20.110  < 2e-16 ***
## sex_comparisonmale-male    0.58733    0.10385   5.656 1.55e-08 ***
## ---
## Signif. codes:  0 '***' 0.001 '**' 0.01 '*' 0.05 '.' 0.1 ' ' 1
## 
## Correlation of Fixed Effects:
##             (Intr) ovrlp_ sx_cmprsnf-
## overlap_est -0.211                   
## sx_cmprsnf- -0.258  0.102            
## sx_cmprsnm- -0.234  0.171  0.792
```

---

### **Correlated Movement**

To evaluate if pairs of giant anteaters exhibited any correlation in
their movement, all dyads with a proximity ratio that differed
significantly from 1 (based on the 95% confidence interval) were
identified and carried forward to our subsequent, correlated movement
analysis.

#### Exploring deviations in proximity ratios

```
#identify pairs that did not have a proximity ratio of 1
proximity_above1 <- proximity_df[proximity_df$proximity_low > 1,]
proximity_below1 <- proximity_df[proximity_df$proximity_high < 1,]

#exclude pairs with a home range overlap of 0
proximity_below1[proximity_below1$overlap_est == 0,]
proximity_below1 <- proximity_below1[proximity_below1$overlap_est != 0,]

#create a dataframe of the deviated pairs
proximity_identified_pairs_df <- rbind(proximity_above1, proximity_below1)
proximity_identified_pairs_df$pair_ID_number <- seq(from = 1, to = 12, by = 1)
proximity_identified_pairs_df <- relocate(proximity_identified_pairs_df, pair_ID_number,
                                          .before = anteater_A)

#correct the sex_comparison output to female-male
proximity_identified_pairs_df <- mutate(proximity_identified_pairs_df, sex_comparison = 
                                          case_when(paste(Sex.A, Sex.B) == "Male Male" ~ "male-male",
                                                    paste(Sex.A, Sex.B) == "Female Female" ~ "female-female",
                                                    paste(Sex.A, Sex.B) == "Male Female" ~ "female-male",
                                                    paste(Sex.A, Sex.B) == "Female Male" ~ "female-male"))

#clean up environment
rm(proximity_above1, proximity_below1)
```

#### Deviated pairs results

```
#number of pairs with a deviated proximity ratio based on sex comparison
table(proximity_identified_pairs_df$sex_comparison)
```

```
## 
## female-female   female-male     male-male 
##             3             6             3
```

```
#test for significance in sex, compare model with and without sex as a variable
proximity_test_pairs <- glmer(proximity_est ~ sex_comparison + (1|site), 
                              family = Gamma(link = "log"), 
                              data = proximity_identified_pairs_df)
```

```
## boundary (singular) fit: see help('isSingular')
```

```
proximity_test2_pairs <- glmer(proximity_est ~ 1 + (1|site), 
                               family = Gamma(link = "log"), 
                               data = proximity_identified_pairs_df)
```

```
## boundary (singular) fit: see help('isSingular')
```

```
proximity_test_results_pairs <- anova(proximity_test_pairs, proximity_test2_pairs)
proximity_test_pvalue_pairs <- round(proximity_test_results_pairs$`Pr(>Chisq)`[2], 2)

#test for significance in home range overlap, compare model with and without overlap as a variable
prox_overlap_test_pairs <- glmer(proximity_est ~ overlap_est + (1|site), 
                                 family = Gamma(link = "log"), 
                                 data = proximity_identified_pairs_df)
```

```
## boundary (singular) fit: see help('isSingular')
```

```
prox_overlap_test2_pairs <- glmer(proximity_est ~ 1 + (1|site), 
                                  family = Gamma(link = "log"), 
                                  data = proximity_identified_pairs_df)
```

```
## boundary (singular) fit: see help('isSingular')
```

```
prox_overlap_test_results_pairs <- anova(prox_overlap_test_pairs, prox_overlap_test2_pairs)
prox_overlap_test_pvalue_pairs <- round(prox_overlap_test_results_pairs$`Pr(>Chisq)`[2], 2)
```

#### Estimating distances in the deviated pairs

```
#subset telemetry data and fitted model for each pair
pair1 <- DATA_TELEMETRY[c("Kyle","Christoffer")]
FIT_pair1 <- FIT[c("Kyle","Christoffer")]

pair2 <- DATA_TELEMETRY[c("Elaine","Christoffer")]
FIT_pair2 <- FIT[c("Elaine","Christoffer")]

pair3 <- DATA_TELEMETRY[c("Kyle","Bumpus")]
FIT_pair3 <- FIT[c("Kyle","Bumpus")]

pair4 <- DATA_TELEMETRY[c("Little_Rick","Elaine")]
FIT_pair4 <- FIT[c("Little_Rick","Elaine")]

pair5 <- DATA_TELEMETRY[c("Makao","Bumpus")]
FIT_pair5 <- FIT[c("Makao","Bumpus")]

pair6 <- DATA_TELEMETRY[c("Puji","Bumpus")]
FIT_pair6 <- FIT[c("Puji","Bumpus")]

pair7 <- DATA_TELEMETRY[c("Rodolfo", "Elaine")]
FIT_pair7 <- FIT[c("Rodolfo", "Elaine")]

pair8 <- DATA_TELEMETRY[c("Larry","Annie")]
FIT_pair8 <- FIT[c("Larry","Annie")]

pair9 <- DATA_TELEMETRY[c("Reid","Larry")]
FIT_pair9 <- FIT[c("Reid","Larry")]

pair10 <- DATA_TELEMETRY[c("Sheron","Maria")]
FIT_pair10 <- FIT[c("Sheron","Maria")]

pair11 <- DATA_TELEMETRY[c("Thomas","Margaret")]
FIT_pair11 <- FIT[c("Thomas","Margaret")]

pair12 <- DATA_TELEMETRY[c("Thomas","Reid")]
FIT_pair12 <- FIT[c("Thomas","Reid")]

#calculate the instantaneous Euclidean distance between each deviated pairs
distance_pair1 <- distances(pair1, FIT_pair1) 
distance_pair2 <- distances(pair2, FIT_pair2)
distance_pair3 <- distances(pair3, FIT_pair3)
distance_pair4 <- distances(pair4, FIT_pair4)
distance_pair5 <- distances(pair5, FIT_pair5)
distance_pair6 <- distances(pair5, FIT_pair5) 
distance_pair7 <- distances(pair7, FIT_pair7)
distance_pair8 <- distances(pair8, FIT_pair8)
distance_pair9 <- distances(pair9, FIT_pair9) 
distance_pair10 <- distances(pair10, FIT_pair10)
distance_pair11 <- distances(pair11, FIT_pair11)
distance_pair12 <- distances(pair12, FIT_pair12)
```

```
#add columns and reorganize dataframe
distance_pair1$pair_ID_number <- 1
distance_pair1$anteater_A <- "Kyle"
distance_pair1$anteater_B <- "Christoffer"
distance_pair1$pair_ID <- paste(distance_pair1$anteater_A, distance_pair1$anteater_B, sep = "_")
distance_pair1 <- relocate(distance_pair1, c(pair_ID_number, pair_ID, anteater_A, anteater_B), .before = low)

distance_pair2$pair_ID_number <- 2
distance_pair2$anteater_A <- "Elaine"
distance_pair2$anteater_B <- "Christoffer"
distance_pair2$pair_ID <- paste(distance_pair2$anteater_A, distance_pair2$anteater_B, sep = "_")
distance_pair2 <- relocate(distance_pair2, c(pair_ID_number, pair_ID, anteater_A, anteater_B), .before = low)

distance_pair3$pair_ID_number <- 3
distance_pair3$anteater_A <- "Kyle"
distance_pair3$anteater_B <- "Bumpus"
distance_pair3$pair_ID <- paste(distance_pair3$anteater_A, distance_pair3$anteater_B, sep = "_")
distance_pair3 <- relocate(distance_pair3, c(pair_ID_number, pair_ID, anteater_A, anteater_B), .before = low)

distance_pair4$pair_ID_number <- 4
distance_pair4$anteater_A <- "Little_Rick"
distance_pair4$anteater_B <- "Elaine"
distance_pair4$pair_ID <- paste(distance_pair4$anteater_A, distance_pair4$anteater_B, sep = "_")
distance_pair4 <- relocate(distance_pair4, c(pair_ID_number, pair_ID, anteater_A, anteater_B), .before = low)

distance_pair5$pair_ID_number <- 5
distance_pair5$anteater_A <- "Makao"
distance_pair5$anteater_B <- "Bumpus"
distance_pair5$pair_ID <- paste(distance_pair5$anteater_A, distance_pair5$anteater_B, sep = "_")
distance_pair5 <- relocate(distance_pair5, c(pair_ID_number, pair_ID, anteater_A, anteater_B), .before = low)

distance_pair6$pair_ID_number <- 6
distance_pair6$anteater_A <- "Puji"
distance_pair6$anteater_B <- "Bumpus"
distance_pair6$pair_ID <- paste(distance_pair6$anteater_A, distance_pair6$anteater_B, sep = "_")
distance_pair6 <- relocate(distance_pair6, c(pair_ID_number, pair_ID, anteater_A, anteater_B), .before = low)

distance_pair7$pair_ID_number <- 7
distance_pair7$anteater_A <- "Rodolfo"
distance_pair7$anteater_B <- "Elaine"
distance_pair7$pair_ID <- paste(distance_pair7$anteater_A, distance_pair7$anteater_B, sep = "_")
distance_pair7 <- relocate(distance_pair7, c(pair_ID_number, pair_ID, anteater_A, anteater_B), .before = low)

distance_pair8$pair_ID_number <- 8
distance_pair8$anteater_A <- "Larry"
distance_pair8$anteater_B <- "Annie"
distance_pair8$pair_ID <- paste(distance_pair8$anteater_A, distance_pair8$anteater_B, sep = "_")
distance_pair8 <- relocate(distance_pair8, c(pair_ID_number, pair_ID, anteater_A, anteater_B), .before = low)

distance_pair9$pair_ID_number <- 9
distance_pair9$anteater_A <- "Reid"
distance_pair9$anteater_B <- "Larry"
distance_pair9$pair_ID <- paste(distance_pair9$anteater_A, distance_pair9$anteater_B, sep = "_")
distance_pair9 <- relocate(distance_pair9, c(pair_ID_number, pair_ID, anteater_A, anteater_B), .before = low)

distance_pair10$pair_ID_number <- 10
distance_pair10$anteater_A <- "Sheron"
distance_pair10$anteater_B <- "Maria"
distance_pair10$pair_ID <- paste(distance_pair10$anteater_A, distance_pair10$anteater_B, sep = "_")
distance_pair10 <- relocate(distance_pair10, c(pair_ID_number, pair_ID, anteater_A, anteater_B), .before = low)

distance_pair11$pair_ID_number <- 11
distance_pair11$anteater_A <- "Thomas"
distance_pair11$anteater_B <- "Margaret"
distance_pair11$pair_ID <- paste(distance_pair11$anteater_A, distance_pair11$anteater_B, sep = "_")
distance_pair11 <- relocate(distance_pair11, c(pair_ID_number, pair_ID, anteater_A, anteater_B), .before = low)

distance_pair12$pair_ID_number <- 12
distance_pair12$anteater_A <- "Thomas"
distance_pair12$anteater_B <- "Reid"
distance_pair12$pair_ID <- paste(distance_pair12$anteater_A, distance_pair12$anteater_B, sep = "_")
distance_pair12 <- relocate(distance_pair12, c(pair_ID_number, pair_ID, anteater_A, anteater_B), .before = low)

#combine into a dataframe
distance_pairs_df <- rbind(distance_pair1, distance_pair2, distance_pair3, distance_pair4,
                           distance_pair5, distance_pair6, distance_pair7, distance_pair8,
                           distance_pair9, distance_pair10, distance_pair11, distance_pair12)

#clean up environment
rm(distance_pair1, distance_pair2, distance_pair3, distance_pair4, distance_pair5,
   distance_pair6, distance_pair7, distance_pair8, distance_pair9, distance_pair10,
   distance_pair11, distance_pair12)
```

```
#subset telemetry data for each individual with a deviated proximity ratio
Bumpus <- DATA_TELEMETRY$Bumpus
Christoffer <- DATA_TELEMETRY$Christoffer
Elaine <- DATA_TELEMETRY$Elaine
Kyle <- DATA_TELEMETRY$Kyle
Little_rick <- DATA_TELEMETRY$Little_Rick
Makao <- DATA_TELEMETRY$Makao
Puji <- DATA_TELEMETRY$Puji
Rodolfo <- DATA_TELEMETRY$Rodolfo
Annie <- DATA_TELEMETRY$Annie
Larry <- DATA_TELEMETRY$Larry
Margaret <- DATA_TELEMETRY$Margaret
Maria <- DATA_TELEMETRY$Maria
Sheron <- DATA_TELEMETRY$Sheron
Reid <- DATA_TELEMETRY$Reid
Thomas <- DATA_TELEMETRY$Thomas

#create a dataframe of an individual's GPS coordinates, format dataset for corrMove analysis
Bumpus_GPS <- data.frame(timestamp = round_date(Bumpus$timestamp, "20 minutes"),
                         Bumpus.x = Bumpus$longitude,
                         Bumpus.y = Bumpus$latitude)
Christoffer_GPS <- data.frame(timestamp = round_date(Christoffer$timestamp, "20 minutes"),
                              Christoffer.x = Christoffer$longitude,
                              Christoffer.y = Christoffer$latitude)
Elaine_GPS <- data.frame(timestamp = round_date(Elaine$timestamp, "20 minutes"),
                         Elaine.x = Elaine$longitude,
                         Elaine.y = Elaine$latitude)
Kyle_GPS <- data.frame(timestamp = round_date(Kyle$timestamp, "20 minutes"),
                       Kyle.x = Kyle$longitude,
                       Kyle.y = Kyle$latitude)
Little_rick_GPS <- data.frame(timestamp = round_date(Little_rick$timestamp, "20 minutes"),
                              Little_rick.x = Little_rick$longitude,
                              Little_rick.y = Little_rick$latitude)
Makao_GPS <- data.frame(timestamp = round_date(Makao$timestamp, "20 minutes"),
                        Makao.x = Makao$longitude,
                        Makao.y = Makao$latitude)
Puji_GPS <- data.frame(timestamp = round_date(Puji$timestamp, "20 minutes"),
                       Puji.x = Puji$longitude,
                       Puji.y = Puji$latitude)
Rodolfo_GPS <- data.frame(timestamp = round_date(Rodolfo$timestamp, "20 minutes"),
                          Rodolfo.x = Rodolfo$longitude,
                          Rodolfo.y = Rodolfo$latitude)
Annie_GPS <- data.frame(timestamp = round_date(Annie$timestamp, "20 minutes"),
                        Annie.x = Annie$longitude,
                        Annie.y = Annie$latitude)
Larry_GPS <- data.frame(timestamp = round_date(Larry$timestamp, "20 minutes"),
                        Larry.x = Larry$longitude,
                        Larry.y = Larry$latitude)
Margaret_GPS <- data.frame(timestamp = round_date(Margaret$timestamp, "20 minutes"),
                           Margaret.x = Margaret$longitude,
                           Margaret.y = Margaret$latitude)
Maria_GPS <- data.frame(timestamp = round_date(Maria$timestamp, "20 minutes"),
                        Maria.x = Maria$longitude,
                        Maria.y = Maria$latitude)
Sheron_GPS <- data.frame(timestamp = round_date(Sheron$timestamp, "20 minutes"),
                         Sheron.x = Sheron$longitude,
                         Sheron.y = Sheron$latitude)
Reid_GPS <- data.frame(timestamp = round_date(Reid$timestamp, "20 minutes"),
                       Reid.x = Reid$longitude,
                       Reid.y = Reid$latitude)
Thomas_GPS <- data.frame(timestamp = round_date(Thomas$timestamp, "20 minutes"),
                         Thomas.x = Thomas$longitude,
                         Thomas.y = Thomas$latitude)

#combine the GPS coordinates of pairs together, reorganize dataframe needed for corrMove into format required and remove duplicate timestamps
cd_pair1 <- merge(Kyle_GPS, Christoffer_GPS)
cd_pair1 <- cd_pair1[, c(1,2,4,3,5)]
cd_pair1 <- cd_pair1[!duplicated(cd_pair1$timestamp),]

cd_pair2 <- merge(Elaine_GPS, Christoffer_GPS)
cd_pair2 <- cd_pair2[, c(1,2,4,3,5)]
cd_pair2 <- cd_pair2[!duplicated(cd_pair2$timestamp),]

cd_pair3 <- merge(Kyle_GPS, Bumpus_GPS, )
cd_pair3 <- cd_pair3[, c(1,2,4,3,5)]
cd_pair3 <- cd_pair3[!duplicated(cd_pair3$timestamp),]

cd_pair4 <- merge(Little_rick_GPS, Elaine_GPS)
cd_pair4 <- cd_pair4[, c(1,2,4,3,5)]
cd_pair4 <- cd_pair4[!duplicated(cd_pair4$timestamp),]

cd_pair5 <- merge(Makao_GPS, Bumpus_GPS)
cd_pair5 <- cd_pair5[, c(1,2,4,3,5)]
cd_pair5 <- cd_pair5[!duplicated(cd_pair5$timestamp),]

cd_pair6 <- merge(Puji_GPS, Bumpus_GPS)
cd_pair6 <- cd_pair6[, c(1,2,4,3,5)]
cd_pair6 <- cd_pair6[!duplicated(cd_pair6$timestamp),]

cd_pair7 <- merge(Rodolfo_GPS, Elaine_GPS)
cd_pair7 <- cd_pair7[, c(1,2,4,3,5)]
cd_pair7 <- cd_pair7[!duplicated(cd_pair7$timestamp),]

cd_pair8 <- merge(Larry_GPS, Annie_GPS)
cd_pair8 <- cd_pair8[, c(1,2,4,3,5)]
cd_pair8 <- cd_pair8[!duplicated(cd_pair8$timestamp),]

cd_pair9 <- merge(Reid_GPS, Larry_GPS)
cd_pair9 <- cd_pair9[, c(1,2,4,3,5)]
cd_pair9 <- cd_pair9[!duplicated(cd_pair9$timestamp),]

cd_pair10 <- merge(Sheron_GPS, Maria_GPS)
cd_pair10 <- cd_pair10[, c(1,2,4,3,5)]
cd_pair10 <- cd_pair10[!duplicated(cd_pair10$timestamp),]

cd_pair11 <- merge(Thomas_GPS, Margaret_GPS)
cd_pair11 <- cd_pair11[, c(1,2,4,3,5)]
cd_pair11 <- cd_pair11[!duplicated(cd_pair11$timestamp),]

cd_pair12 <- merge(Thomas_GPS, Reid_GPS)
cd_pair12 <- cd_pair12[, c(1,2,4,3,5)]
cd_pair12 <- cd_pair12[!duplicated(cd_pair12$timestamp),]
```

#### Estimating correlated movement

For these 12 dyads, we used the methods implemented in the corrMove
package to estimate the amount of correlation in the drift and diffusion
components of those dyads’ movement (Calabrese et al., 2018). Under an
assumption of Brownian motion, the corrMove algorithm estimates
transition points between a family of four possible movement models;
uncorrelated drift and uncorrelated diffusion (UU), correlated drift and
uncorrelated diffusion (CU), uncorrelated drift and correlated diffusion
(UC), and correlated drift and correlated diffusion (CC) (Calabrese et
al., 2018). The most appropriate model for any time window was
identified via AICc based model selection.

```
#Estimate the partition points
prts_pair1 <- findPrts(cd_pair1, W=5, IC = 2)
#Get the MCI estimates and selected model conditional on the data and partition points
cm_pair1 <- corrMove(cd_pair1, prts_pair1)

prts_pair2 <- findPrts(cd_pair2, W=5, IC = 2)
cm_pair2 <- corrMove(cd_pair2, prts_pair2)

prts_pair3 <- findPrts(cd_pair3, W=5, IC = 2)
cm_pair3 <- corrMove(cd_pair3, prts_pair3)

prts_pair4 <- findPrts(cd_pair4, W=5, IC = 2)
cm_pair4 <- corrMove(cd_pair4, prts_pair4)

prts_pair5 <- findPrts(cd_pair5, W=5, IC = 2)
cm_pair5 <- corrMove(cd_pair5, prts_pair5)

prts_pair6 <- findPrts(cd_pair6, W=5, IC = 2)
cm_pair6 <- corrMove(cd_pair6, prts_pair6)

prts_pair7 <- findPrts(cd_pair7, W=5, IC = 2)
cm_pair7 <- corrMove(cd_pair7, prts_pair7)

prts_pair8 <- findPrts(cd_pair8, W=5, IC = 2)
cm_pair8 <- corrMove(cd_pair8, prts_pair8)

prts_pair9 <- findPrts(cd_pair9, W=5, IC = 2)
cm_pair9 <- corrMove(cd_pair9, prts_pair9)

prts_pair10 <- findPrts(cd_pair10, W=5, IC = 2)
cm_pair10 <- corrMove(cd_pair10, prts_pair10)

prts_pair11 <- findPrts(cd_pair11, W=5, IC = 2)
cm_pair11 <- corrMove(cd_pair11, prts_pair11)

prts_pair12 <- findPrts(cd_pair12, W=5, IC = 2)
cm_pair12 <- corrMove(cd_pair12, prts_pair12)
```

#### Correlated movement results

From these analyses, we obtained information on the length of time
for which the dyad exhibited behaviour consistent with the selected
movement model, as well as the correlation coefficients for the drift
and diffusion terms (ranging between –1, indicating avoidance, and 1,
indicating following).

```
#create a new dataframe for correlated movement data
corrmove_df <- proximity_identified_pairs_df

#create empty columns for the results to be saved to
corrmove_df$mean_etaTot.CI.Low <- NA
corrmove_df$mean_etaTot.MLE <- NA
corrmove_df$mean_etaTot.CI.Upp <- NA

#calculate mean total correlated movement
for (i in 1:12) {
  #get the corresponding cm_pair
  cm_pair <- get(paste0("cm_pair", i))
  
  # calculate means
  mean_etaTot_CI_Low <- mean(cm_pair$etaTot.CI.Low)
  mean_etaTot_MLE <- mean(cm_pair$etaTot.MLE)
  mean_etaTot_CI_Upp <- mean(cm_pair$etaTot.CI.Upp)
  
  #save results with mean values to the empty columns
  corrmove_df$mean_etaTot.CI.Low[corrmove_df$pair_ID_number == i] <- mean_etaTot_CI_Low
  corrmove_df$mean_etaTot.MLE[corrmove_df$pair_ID_number == i] <- mean_etaTot_MLE
  corrmove_df$mean_etaTot.CI.Upp[corrmove_df$pair_ID_number == i] <- mean_etaTot_CI_Upp
}

#mean amount of total correlation in all identified pairs movement
round(mean(corrmove_df$mean_etaTot.CI.Low[-1]), 2)
```

```
## [1] 0
```

```
round(mean(corrmove_df$mean_etaTot.MLE[-1]), 2)
```

```
## [1] 0.01
```

```
round(mean(corrmove_df$mean_etaTot.CI.Upp[-1]), 2)
```

```
## [1] 0.03
```

```
#create empty columns for the results to be saved to
corrmove_df$mean_etaDif.MLE <- NA
corrmove_df$mean_etaDft.MLE <- NA

#calculate mean total drift and mean total diffusion correlated movement
for (i in 1:12) {
  # get the corresponding cm_pair
  cm_pair <- get(paste0("cm_pair", i))
  
  # calculate means
  mean_etaDif_MLE <- mean(cm_pair$etaDif.MLE)
  mean_etaDft_MLE <- mean(cm_pair$etaDft.MLE)
  
  #save results with mean values
  corrmove_df$mean_etaDif.MLE[corrmove_df$pair_ID_number == i] <- mean_etaDif_MLE
  corrmove_df$mean_etaDft.MLE[corrmove_df$pair_ID_number == i] <- mean_etaDft_MLE
}

#mean total drift and mean total diffusion correlated movement
round(mean(corrmove_df$mean_etaDif.MLE[-1]), 2)
```

```
## [1] 0.01
```

```
round(mean(corrmove_df$mean_etaDft.MLE[-1]), 2)
```

```
## [1] 0
```

---

#### Case study

As a case example for these general trends, we present a more
in-depth exploration for “Thomas” and “Margaret”, a female-male dyad
that exhibited substantial home-range overlap as well as a proximity
ratio which suggested significant attraction. While we only presented
results for a single dyad, similar patterns were observed across all
dyads.

```
#subset AKDE for each individual
AKDE_thomas <- AKDE["Thomas"]
AKDE_margaret <- AKDE["Margaret"]

#calculate mean home range sizes for Thomas
meta(AKDE_thomas)
```

```
##                  ΔAICc
## Dirac-δ              0
## inverse-Gaussian   Inf
```

```
##                  low      est     high
## mean (km²)  2.339216 2.645406 2.970154
## CoV² (RVAR) 0.000000 0.000000      Inf
## CoV  (RSTD) 0.000000 0.000000      Inf
```

```
#calculate mean home range sizes for Margaret
meta(AKDE_margaret)
```

```
##                  ΔAICc
## Dirac-δ              0
## inverse-Gaussian   Inf
```

```
##                low      est     high
## mean (km²)  2.0671 2.361801 2.675854
## CoV² (RVAR) 0.0000 0.000000      Inf
## CoV  (RSTD) 0.0000 0.000000      Inf
```

```
#calculate home range overlap
round(proximity_identified_pairs_df[11,]$overlap_low, 2)
```

```
## [1] 0.88
```

```
round(proximity_identified_pairs_df[11,]$overlap_est, 2)
```

```
## [1] 0.92
```

```
round(proximity_identified_pairs_df[11,]$overlap_high, 2)
```

```
## [1] 0.94
```

```
#calculate proximity ratio
round(proximity_identified_pairs_df$proximity_low[proximity_identified_pairs_df$pair_ID_number == 11], 2)
```

```
## [1] 0.71
```

```
round(proximity_identified_pairs_df$proximity_est[proximity_identified_pairs_df$pair_ID_number == 11], 2)
```

```
## [1] 0.8
```

```
round(proximity_identified_pairs_df$proximity_high[proximity_identified_pairs_df$pair_ID_number == 11], 2)
```

```
## [1] 0.91
```

```
#calculate distances (measurements are in meters, convert them to km)
round(mean(distance_pairs_df$est[distance_pairs_df$pair_ID_number == 11])/1000, 2)
```

```
## [1] 0.65
```

```
round(min(distance_pairs_df$est[distance_pairs_df$pair_ID_number == 11])/1000, 2)
```

```
## [1] 0
```

```
round(max(distance_pairs_df$est[distance_pairs_df$pair_ID_number == 11])/1000, 2)
```

```
## [1] 2.06
```

```
#calculate the mean 95% CI correlated movement
round(mean(cm_pair11$etaTot.CI.Low), 2)
```

```
## [1] 0.02
```

```
round(mean(cm_pair11$etaTot.MLE), 2)
```

```
## [1] 0.04
```

```
round(mean(cm_pair11$etaTot.CI.Upp), 2)
```

```
## [1] 0.06
```

---

### **Figures**

Recreating figures from the main text.

#### Figure 1

```
# Figure 1: Map

#get bounding box for the geographic location of a city 
bbox <- getbb("Nova Alvorada do Sul Brazil")

#add OSM features for roads
roads <- add_osm_feature(opq(bbox), key = "highway", 
                         value = c("trunk", "motorway", 
                                   "primary", "secondary", "tertiary"))
#retrieve OSM data as an sf object
roads <- osmdata_sf(roads)

#create an overpass query for medium/small streets
secondary_roads <- add_osm_feature(opq(bbox), key = "highway", 
                                   value = c("residential",
                                             "living_street",
                                             "unclassified",
                                             "service",
                                             "footway",
                                             "unpaved",
                                             "motorway_link",
                                             "trunk_link",
                                             "primary_link",
                                             "secondary_link",
                                             "tertiary_link"))
#retrieve OSM data as an sf object
secondary_roads <- osmdata_sf(secondary_roads)

#create an overpass query for medium/small streets
dirt_roads <- add_osm_feature(opq(bbox), key = "highway", 
                              value = c("track"))
#retrieve OSM data as an sf object
dirt_roads <- osmdata_sf(dirt_roads)

#create an overpass query for rivers
rivers <- add_osm_feature(opq(bbox), key = "waterway", 
                          value = c("river"))
#retrieve OSM data as an sf object
rivers <- osmdata_sf(rivers)

#create an overpass query for streams
streams <- add_osm_feature(opq(bbox), key = "waterway", 
                           value = c("stream"))
#retrieve OSM data as an sf object
streams <- osmdata_sf(streams)
```

```
#import raster data
pasture <- terra::rast("data/map/pasture.TIF")
pasture[pasture == 0] <- NA
pasture[pasture == 1] <- 1

planted_forest <- terra::rast("data/map/planted_forest.TIF")
planted_forest[planted_forest == 0] <- NA
planted_forest[planted_forest == 1] <- 3

native_forest <- terra::rast("data/map/native_forest.TIF")
native_forest[native_forest == 0] <- NA
native_forest[native_forest == 1] <- 2
```

```
#add biological data to GPS dataframe
GPS_df <- left_join(GPS_df, bio_df, by = "ID")

#convert the GPS data into a sf object
gps_sf <- st_as_sf(GPS_df, coords = c("GPS.Longitude", "GPS.Latitude"), 
                   crs = 4326)

#find the extent of the GPS coordinates
gps_sf_ext <- ext(gps_sf)

#calculate values for the bounding box that are slightly larger than the extent values of the GPS data
-53.79925 - 0.01
```

```
## [1] -53.80925
```

```
-53.474271 + 0.01
```

```
## [1] -53.46427
```

```
-21.772054 - 0.01
```

```
## [1] -21.78205
```

```
-21.08363 + 0.01
```

```
## [1] -21.07363
```

```
figure1_map <-
  ggplot() +
  geom_spatraster(data = native_forest, aes(fill = native_forest), na.rm = T) +
  geom_spatraster(data = planted_forest, aes(fill = planted_forest), na.rm = T) +
  geom_spatraster(data = pasture, aes(fill = pasture), na.rm = T) +
  scale_fill_gradientn(colours = alpha(c("#eddea4", "#84a98c", "#52796f"), 0.7),
                       na.value = "transparent") +
  geom_sf(data = rivers$osm_lines,
          inherit.aes = FALSE,
          color = "#2c7fb8",
          lwd = 0.3) +
  geom_sf(data = streams$osm_lines,
          inherit.aes = FALSE,
          color = "#2c7fb8",
          lwd = 0.2) +
  geom_sf(data = roads$osm_lines,
          inherit.aes = FALSE,
          color = "#212529",
          lwd = 0.3) +
  geom_sf(data = secondary_roads$osm_lines,
          inherit.aes = FALSE,
          color = "#6c757d",
          lwd = 0.2) +
  geom_sf(data = dirt_roads$osm_lines,
          inherit.aes = FALSE,
          color = "#a68a64",
          lwd = 0.2) +
  geom_sf(data = gps_sf, aes(color = Sex),
          size = 0.05, alpha = 0.5, pch = 16) +
  scale_color_manual(values = c('#004488', '#A50026'), breaks = c('Male', 'Female')) +
  theme_bw() +
  theme(panel.grid.major = element_blank(),
        panel.grid.minor = element_blank(),
        plot.title = element_text(size = 14, family = "sans", face = "bold"),
        axis.title.y = element_text(size=10, family = "sans", face = "bold"),
        axis.title.x = element_text(size=10, family = "sans", face = "bold"),
        axis.text.y = element_text(size=5, family = "sans"),
        axis.text.x  = element_text(size=5, family = "sans"),
        legend.position="none",
        panel.background = element_rect(fill = "#e9ecef"),
        plot.background = element_rect(fill = "transparent", color = NA),
        plot.margin = unit(c(0.2,0.1,0.2,0.2), "cm")) +
  coord_sf(xlim = c(-53.80925, -53.46427),
           ylim = c(-21.78205, -21.07363))
```

#### Figure 2

```
# Figure 2A: Home range size

mean_HR_est <- round(mean(HR_size$HR_est), 2)

figure2a_HR_size <- 
  ggplot() +
  geom_vline(data = HR_size, aes(xintercept = mean_HR_est),
             linetype = "dotdash") +
  geom_linerange(data = HR_size, 
                 aes(xmin = HR_low, xmax = HR_high, y = ID, color = Sex),
                 linewidth = 3, alpha = 0.5) + 
  labs(x = bquote(bold("Home range area" ~ (km^2))),
       y = "") +
  ggtitle("A") +
  scale_color_manual(values = c('#004488', '#A50026'), breaks = c('Male', 'Female')) +
  geom_point(data = HR_size,
             aes(x = HR_est, y = ID, fill = "white"), color = "white",
             size = 2) +
  geom_point(data = HR_size, 
             aes(x = HR_est, y = ID, color = Sex),
             size = 1.5) +
  theme_bw() +
  theme(panel.grid.major = element_blank(),
        panel.grid.minor = element_blank(),
        plot.title = element_text(hjust = -0.1, size = 14, family = "sans", face = "bold"),
        axis.title.y = element_text(size=10, family = "sans", face = "bold"),
        axis.title.x = element_text(size=10, family = "sans", face = "bold"),
        axis.text.y = element_text(size=8, family = "sans"),
        axis.text.x  = element_text(size=8, family = "sans"),
        legend.position="none",
        panel.background = element_rect(fill = "transparent"),
        plot.background = element_rect(fill = "transparent", color = NA))
```

```
# Figure 2B: Home range overlap sex comparison

figure2b_overlap_sex <-
  ggplot(data = overlap_df, 
         mapping = aes(x = sex_comparison, y = overlap_est, fill = sex_comparison)) + 
  geom_boxplot(alpha = 0.5, size = 0.3, outlier.size = 0.3) +
  ylab("Home range overlap") +
  xlab("Sex") +
  ggtitle("B") +
  theme_bw() +
  theme(panel.grid.major = element_blank(),
        panel.grid.minor = element_blank(),
        plot.title = element_text(size = 14, family = "sans", face = "bold"),
        axis.title.y = element_text(size=10, family = "sans", face = "bold"),
        axis.title.x = element_blank(),
        axis.text.y = element_text(size=8, family = "sans"),
        axis.text.x  = element_text(size=8, family = "sans", face = "bold"),
        legend.position="none",
        panel.background = element_rect(fill = "transparent"),
        plot.background = element_rect(fill = "transparent", color = NA)) +
  scale_fill_manual(values = c("#A50026", "#9970AB", "#004488"),
                    breaks = c("female-female","female-male","male-male"),
                    labels = c("Female - Female", "Female - Male", "Male - Male")) +
  scale_y_continuous(limits = c(0,1)) +
  scale_x_discrete(breaks = c("female-female","female-male","male-male"),
                   labels = c("Female - Female", "Female - Male", "Male - Male"))
```

```
#multi-panel
figure2_left <- grid.arrange(figure2a_HR_size,
                             figure2b_overlap_sex,
                             nrow = 2, heights = c(0.65, 0.35))
```

```
# Figure 2 C,D: Home range overlap

#assign colours based on sex of individual at site 1
COL_1 <- c("#004488", "#004488", "#A50026", "#A50026", "#004488", 
           "#A50026", "#004488", "#004488", "#004488", "#A50026", 
           "#A50026", "#004488")

#assign colours based on sex of individual at site 2
COL_2 <- c("#A50026", "#004488", "#A50026", "#A50026", "#004488", 
           "#004488", "#A50026", "#A50026", "#004488", "#A50026", 
           "#004488") 

png(file = "figures/individual figures/figure2_right_overlap.png", 
    width = 2.86, height = 6, units = "in", res = 600)
par(mfrow=c(2,1))
par(mgp = c(2, 0.5, 0))             
par(mar = c(3, 3, 1.25, 0.25))      
plot(AKDE_1, 
     col.DF = COL_1, 
     col.level = COL_1, 
     col.grid = NA, 
     level = NA,
     lwd.level = 1,            
     cex.lab = 1,              
     cex.axis = 0.8,           
     font.lab = 2)             
title("C", adj = 0)
plot(AKDE_2, 
     col.DF = COL_2, 
     col.level = COL_2, 
     col.grid = NA, 
     level = NA,
     lwd.level = 1,            
     cex.lab = 1,              
     cex.axis = 0.8,           
     font.lab = 2)             
title("D", adj = 0)
dev.off()
```

#### Figure 3

```
# Figure 3A: Proximity ratio

figure3a_proximity_ratio <- 
  ggplot() +
  geom_hline(yintercept = 1, col = "grey70", linetype = "dashed") +
  geom_point(data = proximity_df, 
             aes(y = proximity_est, x = overlap_est, col = sex_comparison),
             size = 1.2, alpha = 0.3, shape = 16) + 
  geom_segment(data = proximity_df, 
               aes(x = overlap_est, xend = overlap_est, 
                   y = proximity_low, yend = proximity_high, 
                   col = sex_comparison), 
               linewidth = 0.3, alpha = 0.3) +
  geom_point(data = proximity_identified_pairs_df, 
             aes(y = proximity_est, x = overlap_est, col = sex_comparison),
             size = 1.2) +
  geom_segment(data = proximity_identified_pairs_df,
               aes(x = overlap_est, xend = overlap_est, 
                   y = proximity_low, yend = proximity_high, 
                   col = sex_comparison), 
               linewidth = 0.3) +
  scale_x_continuous(limits = c(0,1), expand = c(0,0.02)) +
  scale_color_manual(values = c("#A50026", "#9970AB", "#004488"),
                     labels = c("Female - Female", "Female - Male", "Male - Male"),
                     name = "") +
  ylab("Proximity ratio") +
  xlab("Home-range overlap") +
  ggtitle("A") +
  theme_bw() +
  theme(panel.grid.major = element_blank(),
        panel.grid.minor = element_blank(),
        plot.title = element_text(hjust = 0.005, size = 14, family = "sans", face = "bold"),
        axis.title.y = element_text(size=10, family = "sans", face = "bold"),
        axis.title.x = element_text(size=10, family = "sans", face = "bold"),
        axis.text.y = element_text(size=8, family = "sans"),
        axis.text.x  = element_text(size=8, family = "sans"),
        legend.text = element_text(size=6, family = "sans", face = "bold"),
        legend.position = c(0.8, 0.9),
        legend.key.height = unit(0.3, "cm"),
        legend.key=element_blank(),
        panel.background = element_rect(fill = "transparent"),
        legend.background = element_rect(fill = "transparent"),
        plot.background = element_rect(fill = "transparent", color = NA),
        plot.margin = unit(c(0.2,0.1,0.2,0.2), "cm"))
```

```
# Figure 3B: Encounters

#exclude 0 encounters
proximity_df2 <- proximity_df[proximity_df$encounter_count != 0,]

figure3b_encounters <-
  ggplot(data = proximity_df2,
         aes(y = encounter_count, x = overlap_est)) +
  geom_point(data = proximity_df2, 
             aes(y = encounter_count, x = overlap_est, col = sex_comparison),
             size = 1.2) + 
  geom_smooth(method="lm", formula = y ~ x, se=F, col = "black") +
  scale_y_log10(expand = c(0,0.1), limits = c(0.1,2000),
                breaks = trans_breaks("log10", function(x) 10^x),
                labels = trans_format("log10", math_format(10^.x))) +
  scale_x_continuous(limits = c(0,1), expand = c(0,0.02)) +
  scale_color_manual(values = c("#A50026", "#9970AB", "#004488"),
                     labels = c("Female - Female", "Female - Male", "Male - Male"),
                     name = "") +
  xlab("Home-range overlap") +
  ylab("Encounter count") +
  ggtitle("B") +
  theme_bw() +
  theme(panel.grid.major = element_blank(),
        panel.grid.minor = element_blank(),
        plot.title = element_text(size = 14, family = "sans", face = "bold"),
        axis.title.y = element_text(size=10, family = "sans", face = "bold"),
        axis.title.x = element_text(size=10, family = "sans", face = "bold"),
        axis.text.y = element_text(size=8, family = "sans"),
        axis.text.x  = element_text(size=8, family = "sans"),
        legend.position="none",
        panel.background = element_rect(fill = "transparent"),
        legend.background = element_rect(fill = "transparent"),
        plot.background = element_rect(fill = "transparent", color = NA),
        plot.margin = unit(c(0.2,0.1,0.2,0.2), "cm"))
```

```
# Figure 3C: Encounters over time

#set threshold of 15m
distance_df$encounter <- ifelse(distance_df$distance_est > 15, 0,1)
encounter_df <- distance_df[which(distance_df$encounter == 1),]
encounter_df$doy <- yday(encounter_df$timestamp)
encs <- aggregate(encounter ~ sex_comparison + doy + pair_ID, 
                  data = encounter_df, FUN = "sum")
encs <- merge(encs, proximity_df[,c("pair_ID","sex_comparison", "site")])

figure3c_encounters_overtime <-
  ggplot(data = encs,
         aes(y = encounter, x = doy, col = sex_comparison)) +
  geom_bar(stat = "identity", position = "stack") + 
  scale_x_continuous(limits = c(-2,370), expand = c(0,1)) +
  scale_y_continuous(limits = c(0,110), expand = c(0,1)) +
  scale_color_manual(values = c("#A50026", "#9970AB", "#004488"),
                     labels = c("Female - Female", "Female - Male", "Male - Male"),
                     name = "") +
  xlab("Day of year") +
  ylab("Encounter count") +
  ggtitle("C") +
  theme_bw() +
  theme(panel.grid.major = element_blank(),
        panel.grid.minor = element_blank(),
        plot.title = element_text( size = 14, family = "sans", face = "bold"),
        axis.title.y = element_text(size=10, family = "sans", face = "bold"),
        axis.title.x = element_text(size=10, family = "sans", face = "bold"),
        axis.text.y = element_text(size=8, family = "sans"),
        axis.text.x  = element_text(size=8, family = "sans"),
        legend.position="none",
        panel.background = element_rect(fill = "transparent"),
        legend.background = element_rect(fill = "transparent"),
        plot.background = element_rect(fill = "transparent", color = NA),
        plot.margin = unit(c(0.2,0.1,0.2,0.2), "cm"))
```

```
#multi-panel
figure3_top <- grid.arrange(figure3a_proximity_ratio,
                            figure3b_encounters,
                            ncol = 2)
figure3 <- grid.arrange(figure3_top,
                        figure3c_encounters_overtime,
                        nrow = 2,
                        heights = c(0.35, 0.35))
```

#### Figure 4

```
# Figure 4A: Home range overlap (case study)

#subset home range overlap data
AKDE_pair11 <- AKDE[c("Thomas", "Margaret")]
#assign colours
COL_pair11 <- c("#004488", "#A50026")

#plot case study pair home range overlap with GPS points
png(file = "figures/individual figures/figure4a_overlap_pair11.png", 
    width = 2.86, height = 4, units = "in", res = 600)
par(mgp = c(1.5, 0.5, 0))                         
par(mar = c(2.5, 2.5, 1.75, 0.5))                   
plot(AKDE_pair11, col.DF = COL_pair11, 
     col.level = COL_pair11, 
     col.grid = NA, 
     level = NA,
     lwd.level = 1,            
     cex.lab = 0.9,                 
     cex.axis = 0.7,                 
     font.lab = 2,                  
     tcl  = -0.35)                            
title("A", adj = 0)
plot(list(Thomas, Margaret), 
     error = FALSE,
     col = c("#004488", "#A50026"),
     add = TRUE)
dev.off()
```

```
# Figure 4B: Home range overlap (case study)

#subset distance data
distance_pair11 <- distance_pairs_df[distance_pairs_df$pair_ID_number == 11,]

figure4b_pair11_distance <- 
  ggplot() +
  geom_line(data = distance_pair11,
            aes(y = est, x = timestamp, 
            ), size = 0.15) +
  xlab("") +
  ylab("Distance (m)") +
  ggtitle("B") +
  theme_bw() +
  theme(panel.grid.major = element_blank(),
        panel.grid.minor = element_blank(),
        plot.title = element_text(size = 14, family = "sans", face = "bold"),
        axis.title.y = element_text(size=10, family = "sans", face = "bold"),
        axis.title.x = element_text(size=10, family = "sans", face = "bold"),
        axis.text.y = element_text(size=8, family = "sans"),
        axis.text.x  = element_text(size=8, family = "sans"),
        panel.background = element_rect(fill = "transparent"),
        plot.background = element_rect(fill = "transparent", color = NA),
        plot.margin = unit(c(0.1,0.1,0.05,0.2), "cm"))
```

```
# Figure 4C: Correlated movement (total correlated) 

figure4c_pair11_cm <-
  ggplot() +
  geom_hline(aes(yintercept = 0),
             linetype = "dashed", alpha = 0.7) +
  geom_point(aes(y = cm_pair11$etaTot.MLE, x = cm_pair11$timestamp, 
                 col = cm_pair11$sel.mod.code), 
             size = 0.5) +
  xlab("") +
  ylab("Total correlation") +
  ggtitle("C") +
  scale_color_manual(values = c('#EE7733', '#88CCEE', '#BBBBBB'), 
                     breaks = c('CU', 'UC', 'UU')) + 
  theme_bw() +
  scale_y_continuous(limits = c(-1,1)) +
  theme(panel.grid.major = element_blank(),
        panel.grid.minor = element_blank(),
        plot.title = element_text(size = 14, family = "sans", face = "bold"),
        axis.title.y = element_text(size=10, family = "sans", face = "bold"),
        axis.title.x = element_text(size=10, family = "sans", face = "bold"),
        axis.text.y = element_text(size=8, family = "sans"),
        axis.text.x  = element_text(size=8, family = "sans"),
        legend.direction = "horizontal",
        legend.position = c(0.5,1.1),     
        legend.title = element_blank(),
        legend.background = element_rect(fill = "transparent"),
        panel.background = element_rect(fill = "transparent"),
        plot.background = element_rect(fill = "transparent", color = NA),
        plot.margin = unit(c(0.1,0.1,0.05,0.2), "cm")) + 
  guides(colour = guide_legend(override.aes = list(size=3)))
```

```
figure4_right <- grid.arrange(figure4b_pair11_distance, 
                              figure4c_pair11_cm,
                              ncol=1)
```

---

### End Session

```
sessionInfo()
```

```
## R version 4.3.0 (2023-04-21 ucrt)
## Platform: x86_64-w64-mingw32/x64 (64-bit)
## Running under: Windows 10 x64 (build 19045)
## 
## Matrix products: default
## 
## 
## locale:
## [1] LC_COLLATE=English_Canada.utf8  LC_CTYPE=English_Canada.utf8   
## [3] LC_MONETARY=English_Canada.utf8 LC_NUMERIC=C                   
## [5] LC_TIME=English_Canada.utf8    
## 
## time zone: Asia/Bangkok
## tzcode source: internal
## 
## attached base packages:
## [1] stats     graphics  grDevices utils     datasets  methods   base     
## 
## other attached packages:
##  [1] corrMove_0.1.0  glmmTMB_1.1.7   lme4_1.1-33     Matrix_1.5-4   
##  [5] ctmm_1.2.0      sf_1.0-12       tidyterra_0.4.0 terra_1.7-29   
##  [9] osmdata_0.2.5   scales_1.2.1    gridExtra_2.3   lubridate_1.9.2
## [13] tibble_3.2.1    tidyr_1.3.0     dplyr_1.1.2     ggplot2_3.4.2  
## [17] readr_2.1.4    
## 
## loaded via a namespace (and not attached):
##  [1] gtable_0.3.3        TMB_1.9.4           xfun_0.39          
##  [4] bslib_0.4.2         raster_3.6-20       lattice_0.21-8     
##  [7] tzdb_0.3.0          numDeriv_2016.8-1.1 vctrs_0.6.2        
## [10] tools_4.3.0         generics_0.1.3      parallel_4.3.0     
## [13] proxy_0.4-27        fansi_1.0.4         highr_0.10         
## [16] DEoptimR_1.1-2      pkgconfig_2.0.3     KernSmooth_2.23-20 
## [19] lifecycle_1.0.3     compiler_4.3.0      statmod_1.5.0      
## [22] munsell_0.5.0       codetools_0.2-19    Bessel_0.6-0       
## [25] htmltools_0.5.5     class_7.3-21        sass_0.4.6         
## [28] gmp_0.7-2           yaml_2.3.7          crayon_1.5.2       
## [31] pillar_1.9.0        nloptr_2.0.3        jquerylib_0.1.4    
## [34] MASS_7.3-58.4       classInt_0.4-9      cachem_1.0.8       
## [37] boot_1.3-28.1       robustbase_0.99-0   RSpectra_0.16-1    
## [40] nlme_3.1-162        tidyselect_1.2.0    digest_0.6.31      
## [43] purrr_1.0.1         splines_4.3.0       fastmap_1.1.1      
## [46] grid_4.3.0          expm_0.999-7        colorspace_2.1-0   
## [49] cli_3.6.1           magrittr_2.0.3      utf8_1.2.3         
## [52] e1071_1.7-13        Rmpfr_0.9-3         withr_2.5.0        
## [55] bit64_4.0.5         sp_1.6-0            timechange_0.2.0   
## [58] Gmedian_1.2.7       rmarkdown_2.21      bit_4.0.5          
## [61] hms_1.1.3           evaluate_0.21       knitr_1.42         
## [64] rgdal_1.6-6         rlang_1.1.1         Rcpp_1.0.10        
## [67] glue_1.6.2          DBI_1.1.3           vroom_1.6.3        
## [70] rstudioapi_0.14     minqa_1.2.5         jsonlite_1.8.4     
## [73] R6_2.5.1            units_0.8-2
```

---

### References

Bates, D., Mächler, M., Bolker, B. & Walker, S. (2015). Fitting
Linear Mixed-Effects Models Using lme4. J. Stat. Soft. 67.

Brooks, M., E., Kristensen, K., Benthem, K., J. ,van, Magnusson, A.,
Berg, C., W., Nielsen, A., Skaug, H., J., Mächler, M. & Bolker, B.,
M. (2017). glmmTMB Balances Speed and Flexibility Among Packages for
Zero-inflated Generalized Linear Mixed Modeling. The R Journal 9,
378.

Calabrese, J.M. & Fleming, C.H. (2023). corrMove: Analyze
Correlated Movements in Multi-Individual Datasets.

Calabrese, J.M., Fleming, C.H., Fagan, W.F., Rimmler, M., Kaczensky,
P., Bewick, S., Leimgruber, P. & Mueller, T. (2018). Disentangling
social interactions and environmental drivers in multi-individual
wildlife tracking data. Phil. Trans. R. Soc. B 373, 20170007.

Calabrese, J.M., Fleming, C.H. & Gurarie, E. (2016). ctmm: an R
package for analyzing animal relocation data as a continuous‐time
stochastic process. Methods Ecol Evol 7, 1124–1132.

Fleming, C.H. & Calabrese, J.M. (2017). A new kernel density
estimator for accurate home‐range and species‐range area estimation.
Methods Ecol Evol 8, 571–579.

Fleming, C.H. & Calabrese, J.M. (2022). ctmm: Continuous-Time
Movement Modeling.

Fleming, C.H., Fagan, W.F., Mueller, T., Olson, K.A., Leimgruber, P.
& Calabrese, J.M. (2015). Rigorous home range estimation with
movement data: a new autocorrelated kernel density estimator. Ecology
96, 1182–1188.

Noonan, M.J., Ascensão, F., Yogui, D.R. & Desbiez, A.L.J. (2022).
Roads as ecological traps for giant anteaters. Animal Conservation 25,
182–194.

Noonan, M.J., Tucker, M.A., Fleming, C.H., Akre, T.S., Alberts, S.C.,
Ali, A.H., Altmann, J., Antunes, P.C., Belant, J.L., Beyer, D., Blaum,
N., Böhning‐Gaese, K., Cullen, L., De Paula, R.C., Dekker, J.,
Drescher‐Lehman, J., Farwig, N., Fichtel, C., Fischer, C., Ford, A.T.,
Goheen, J.R., Janssen, R., Jeltsch, F., Kauffman, M., Kappeler, P.M.,
Koch, F., LaPoint, S., Markham, A.C., Medici, E.P., Morato, R.G.,
Nathan, R., Oliveira‐Santos, L.G.R., Olson, K.A., Patterson, B.D.,
Paviolo, A., Ramalho, E.E., Rösner, S., Schabo, D.G., Selva, N.,
Sergiel, A., Xavier Da Silva, M., Spiegel, O., Thompson, P., Ullmann,
W., Zięba, F., Zwijacz‐Kozica, T., Fagan, W.F., Mueller, T. &
Calabrese, J.M. (2019). A comprehensive analysis of autocorrelation and
bias in home range estimation. Ecological Monographs 89.

R Core Team. (2022). R: A language and environment for statistical
computing.

Smithson, M. & Verkuilen, J. (2006). A better lemon squeezer?
Maximum-likelihood regression with beta-distributed dependent variables.
Psychological Methods 11, 54–71.

Wickham, H. (2016). ggplot2: Elegant Graphics for Data Analysis.
Springer-Verlag New York.

Winner, K., Noonan, M.J., Fleming, C.H., Olson, K.A., Mueller, T.,
Sheldon, D. & Calabrese, J.M. (2018). Statistical inference for home
range overlap. Methods Ecol Evol 9, 1679–1691.
